# Detection of Melanoma Using Deep Serum Proteome Profiling and Machine Learning

**DOI:** 10.64898/2026.06.11.731698

**Authors:** David R. Hall, Brianna I. Gonzalez, Athena A. Schepmoes, Thomas L. Fillmore, Tyler J. Sagendorf, Bennett Drucker, Karin D. Rodland, J. Charles A. Lacson, Clifton L. Dalgard, Isaac Brownell, Jerry S.H. Lee, Craig D. Shriver, Elaine S. Keung, Liesl S. Grenier, Vladislav A. Petyuk, Tao Liu

## Abstract

**Background:** Early detection is the primary determinant of melanoma survival, yet screening still relies on visual inspection, and no serum protein biomarkers are in clinical use for early detection. We sought to identify serum protein signatures of melanoma development in a longitudinal military cohort.

**Methods:** Using the Department of Defense Serum Repository, we analyzed 390 serum samples from 73 melanoma cases and 73 matched controls, collected at up to four timepoints spanning 4 years before to 2 years after diagnosis. Samples were profiled by data-independent acquisition mass spectrometry with nanoparticle-based protein enrichment. Proteins were filtered using a signal-to-noise metric derived from pooled quality control samples. Random forest classifiers with leave-one-out cross-validation, univariate prefiltering, and Boruta feature selection were applied to cross-sectional case–control and longitudinal within-subject comparisons. Performance was assessed by the area under the receiver operating characteristic curve, and models were compared using DeLong’s test.

**Results:** We quantified 3,364 proteins. Cross-sectional classification in the diagnosis window (≤1 year post-diagnosis) reached an area under the curve of 0.823 (95% confidence interval, 0.723–0.923), driven by an eight-protein consensus signature (PRSS1, GSK3B, FAM20C, CES1, CCL14, EPHA10, LMAN2, ITIH1) spanning proteases, immune activation, and extracellular matrix remodeling. Discrimination was weak before diagnosis, with areas under the curve of 0.609 at 2 years and 0.566 at 4 years prior, the latter not distinguishable from chance. The serine proteases PRSS1 and PRSS2 recurred across cross-sectional and longitudinal models. Adding age or military service branch did not improve classification beyond proteins alone.

**Conclusions:** Deep serum proteome profiling distinguishes melanoma cases from matched controls in the diagnosis window and yields candidate markers, with weak signals preceding clinical diagnosis. In the absence of external validation, these findings provide preliminary evidence for a minimally invasive, serum protein-based approach to melanoma detection that could complement visual screening and help triage at-risk individuals for dermatologic evaluation, pending validation in larger, more diverse cohorts.

## Background

Cutaneous melanoma is the fifth most common cancer in the United States, with incidence steadily increasing at approximately 1.6% annually and an estimated 112,000 new diagnoses expected in 2026.[1] Melanoma incidence is particularly elevated among active-duty military personnel, representing the most commonly diagnosed non-sex-specific cancer in this population from 2005 to 2014 and accounting for 17.5% of cancer diagnoses.[2] Given that 5-year survival drops from >99% for localized disease to 35% for distant metastases, early detection fundamentally influences outcomes.[1]

Current melanoma detection relies primarily on visual inspection by dermatologists, an approach that is subjective, requires specialized expertise, and cannot scale to population-level screening.[3] Moreover, certain melanoma subtypes, including spitzoid and desmoplastic melanomas, pose particular diagnostic challenges, as they can be difficult to distinguish from their benign counterparts even for experienced dermatopathologists.[4] While substantial progress has been made in understanding melanoma biology through genomic and proteomic analysis of tumor tissue[5–7], these approaches require invasive biopsy and cannot be used for routine monitoring of at-risk populations.[8]

Blood-based biomarkers are reported for advanced melanomas: S100B, lactate dehydrogenase (LDH), and circulating tumor DNA (ctDNA) are used clinically for prognosis and monitoring treatment response.[9–11] However, no serum biomarkers are routinely used for early melanoma detection. While some multi-cancer early detection (MCED) tests include melanoma among their targets, these primarily employ genetic, epigenetic, and fragmentomic assays with limited reported sensitivity for early-stage disease.[12] Protein-based serum biomarkers remain underexplored for melanoma detection, representing an opportunity for complementary approaches where routine laboratory testing could help triage individuals for follow-up dermatologic evaluation.

Recent advances in mass spectrometry-based proteomics offer new possibilities for biomarker discovery. Data-independent acquisition (DIA) mass spectrometry enables comprehensive, untargeted quantification of thousands of proteins simultaneously[13,14], while nanoparticle-based enrichment technologies such as the Seer Proteograph platform expand proteome coverage by compressing the dynamic range of the serum proteome.[15,16] Together, these technologies enable deep serum proteome profiling suitable for discovery of novel protein signatures.

Here, we use the Department of Defense Serum Repository (DODSR),[17] which contains longitudinal serum samples from military personnel collected before and after melanoma diagnosis. Using DIA mass spectrometry with Seer Proteograph nanoparticle enrichment, we analyzed serum samples spanning 4 years before to 2 years after diagnosis. Our analytical approach combines cross-sectional case-control comparisons with longitudinal tracking of within-individual protein changes, enabling identification of candidate protein signatures for potential future validation.

## Methods

### Study sample selection

Melanoma cases were identified using the Department of Defense’s (DoD) Joint Pathology Center’s (JPC) Automated Central Tumor Registry (ACTUR) using International Classification of Diseases for Oncology Third Edition (ICD-O-3) morphology codes 8720-8723, 8762, 8730, 8740-8746, 8761, 8771-8774, site code C43, and behavior code 3. Equivalent ICD-9 and -10 site codes were also used to identify cases. Cases were diagnosed while on active duty from 2001 to 2016 and were stage T2 or higher at diagnosis. Cases with a history of other cancers were excluded, except for non-melanoma skin cancers (NMSC). Controls were matched 1:1 by race and ethnicity, age (±1 yr), sex, and years (±1 yr) of serum sample acquisition (Figure S1). Data on these matching variables and service branch were provided by the Armed Forces Health Surveillance Division. Controls were on active duty during the period of 2001 to 2016. Individuals with a history of any cancer (including NMSC) reported to ACTUR were excluded. Individuals with a history of organ transplants, immunological disorders, bone marrow treatment, bone marrow failure, diseases of white blood cells, blood, and blood forming organs, as identified through ICD-9 and -10 codes, were excluded from the study.

### Pathologic data

The following data were obtained from ACTUR for each case: anatomic site of the primary tumor, clinical stage, pathologic stage, Breslow depth, status of perineural invasion, status of lymphovascular invasion, regression, presence of ulceration/inflammation, and the type of metastases (local, regional, distant). These variables were used for cohort characterization but not incorporated into biomarker modeling, as they are obtained post-biopsy and thus unavailable in the intended screening context.

### DODSR serum samples

Our study design follows the mixed longitudinal biomarker discovery framework we previously established for head and neck cancer detection.[18] DODSR specimens are residual serum from routine HIV screening and preand post-deployment blood draws, separated and frozen at central processing laboratories and stored at −30 °C.[17] Briefly, we requested up to four serum samples per case or control: (1) the diagnosis-window sample (“DX window”), defined as the serum specimen collected closest to clinical melanoma diagnosis (within 1 year after diagnosis), (2) approximately 2 years prior (“DX_2yr_prior”), (3) 4 years prior (“DX_4yr_prior”), and (4) 2 years post-diagnosis (“DX_2yr_post”) when available (Figure 1B). We prioritized samples with complete longitudinal serum samples, but due to deployment schedules and routine collection intervals, not all subjects had complete sampling across all timepoints, resulting in varying sample sizes per timepoint (Figure 1C). This mixed longitudinal study design enabled two complementary analytical approaches: cross-sectional comparisons between cases and controls at matched timepoints, and longitudinal comparisons tracking within-individual protein changes across the disease trajectory (Figure 1D). The incomplete sampling across timepoints necessitated treating each timepoint comparison independently rather than employing traditional repeated-measures longitudinal models. To establish a technical baseline for data quality assessment, external quality control (QC) samples were included throughout the study. These consisted of off-the-clot pooled serum from 20 male donors (12 African American, 8 Caucasian, median age 47 years; SKU: SD1060-P, GoldenWest BioSolutions LLC, Temecula, CA).

**Figure 1:**
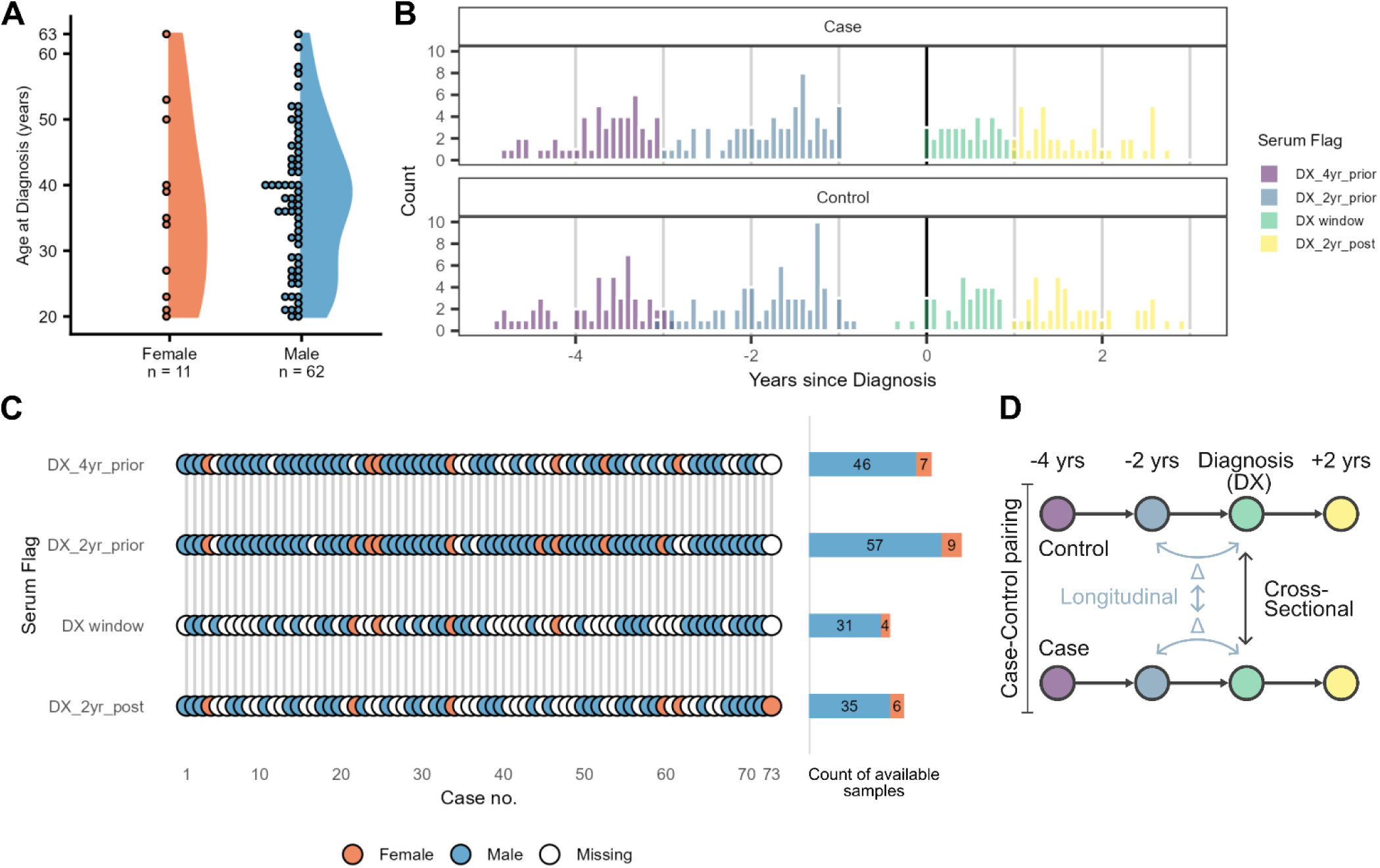
Study cohort summary and design. (A) Age and sex distribution of study subjects at diagnosis. (B) Distribution of serum collection times relative to diagnosis. (C) Sample availability across timepoints for matched case-control pairs. (D) Illustration of cross-sectional (between case-control at each timepoint) and longitudinal (temporal differences within subjects) analysis approaches.

### Preparation of serum samples

Serum samples were processed using the Seer Proteograph XT assay (Seer Inc., Redwood City, CA), a commercially available nanoparticle-based protein enrichment workflow applicable to both serum and plasma.[19] The assay utilizes two distinct pools of engineered nanoparticles (NP-A and NP-B) that differ in surface chemistry and recover largely overlapping sets of proteins with different efficiencies (pooled plasma, Figure S2).[15,16,20,21] Corona formation compresses the dynamic range of serum, so measured intensities reflect relative rather than native abundance. During operation, each serum specimen is split and processed with both nanoparticle pools, yielding 80 peptide aliquots from 40 serum samples in one batch. Seer sample batching followed a structured randomization protocol: (1) longitudinal specimens from matched case-control pairs (ranging from 1 to 4 timepoints per pair) were assigned to the same batch; (2) each batch included 4 external QC samples sourced from the same commercial pooled serum; and (3) a total of 11 batches were required to accommodate all study samples and QCs (Figure S3, Supplementary Information). Case-control pairs were randomly assigned to batches, after which samples within each batch were randomly distributed across the 40 available processing positions.

On the day of processing, samples were thawed on ice. Serum aliquots of 220 μL were brought to the required 250 μL volume using the proprietary Seer assay buffer.[22] The Proteograph XT workflow was performed according to manufacturer’s instructions, including automated trypsin digestion. Post-digestion peptide yield was quantified using the Pierce Fluorescence Quantitative Assay (Thermo Fisher Scientific, Waltham, MA). Samples were dried under vacuum and stored at -80 °C until analysis using liquid chromatography coupled to mass spectrometry (LC-MS) in the DIA mode.

### LC-MS Data-Independent Acquisition (DIA)

Samples were analyzed via DIA in the same batch order as their Seer preparation. Prior to LC-MS acquisition samples were transferred to Evotips using a semi-automated workflow. First, peptides were reconstituted to 25 ng/µl in 3% ACN, 0.1% FA in H_2_O using the Seer SP100 liquid handling robot. Then, using an Opentrons OT-2 (Opentrons Labworks Inc., Long Island City, NY) equal volumes of NP-A and NP-B peptides from the same serum specimen were combined and spiked with 1× iRT peptides (Biognosys AG, Zurich Switzerland). Combined peptides[23] were then loaded onto Evotips using the OT-2 liquid handling robot.[24] Given the substantial overlap in protein groups detected by NP-A and NP-B, equal masses were combined to reduce the number of LC-MS acquisitions while maintaining proteome coverage (Figure S2). Thus, each LC-MS injection consists of 500 ng of peptides (250 ng NP-A and 250 ng NP-B). The quantitative accuracy of this workflow was verified using a mixed-species experiment,[25,26] which demonstrated reliable measurement of protein fold changes ranging from 2-fold to 100-fold (Figure S4).

Samples were run on an Evosep One system[27] on a 15 cm × 150 µm inner diameter (i.d.) column packed with 1.5 µm C18 beads by Dr. Maisch (EV1137) maintained at 50 °C and interfaced to a 30 µm i.d. stainless steel emitter and separated using the 15 samples per day (SPD) extended gradient method (all from Evosep Inc., Odense Denmark). MS acquisition was performed on a Thermo Fisher Exploris 480 quadrupole-orbitrap hybrid mass spectrometer (Thermo Fisher Scientific, Waltham, MA) operating in DIA mode in ESI positive in profile mode, and a default charge state of 3. The MS1 scan was between 400 to 1000 m/z at a resolution of 120,000 with a Normalized AGC target of 300%. Subsequently, 30 variable- width MS2 scans covering 400 to 1000 m/z were performed at a resolution of 30,000 and a normalized AGC target of 1000%. Both external quality control samples and XIC traces of iRT peptides were monitored throughout data acquisition to ensure instrument operations were within acceptable boundaries.[28]

### MS/MS searches

Post-acquisition, .raw files were converted to .htrms (HTRMS Converter 19.7, Biognosys AG) and searched using Spectronaut (19.7.250203, Biognosys AG)[29] in DirectDIA+ mode against a custom FASTA database comprising GENCODE release 42 and Ensembl release 108 sequences supplemented with 203 common LC-MS proteomics contaminants, totaling 51,075 unique entries.[30] DirectDIA+ search settings were kept as default with a precursor Q-Value cutoff < 0.01; Precursor PEP cutoff < 0.2; Protein Q-Value Cutoff (Experiment) < 0.01; Protein Q-Value Cutoff (Run) < 0.05. Protein groups identified from a single peptide were retained[31]; the reproducibility of their quantification was assessed downstream by signal-tonoise and presence filtering. Protein group results were exported using the BGS Factory Report (Normal) scheme for downstream analysis. Post-search, Ensembl protein annotations were extended to UniProt IDs[32] using biomaRt[33] for enhanced functional annotation.

### Data preprocessing and statistical analysis

Protein group normalization was performed using Spectronaut’s default normalization parameters prior to exporting data.[34] Exported protein group results were analyzed in R (4.5.1). Data were first assembled into an MSnSet object and the non-keratin related contaminants were removed.[35] Subsequently protein group intensities were log2 transformed and batch corrected using ComBat.[36] Owing to distinct variance patterns between external QC pooled serum and DODSR study specimens, datasets were separated into QC-only and study-only subsets, batch corrected separately using ComBat and subsequently recombined for analysis (Figure S5 and S6).

The signal-to-noise ratio for each protein was calculated as the ratio of variance across DODSR samples to variance across external QC samples, see Eq. 1.[18] Since DODSR sample variance reflects both biological and technical variation while QC variance represents technical variation alone, this ratio provides an estimate of signal-to-noise for each protein. Consistent with our prior biomarker discovery work,[18] proteins with S/N ≥ 2 were retained for downstream analysis. This threshold identified 1,807 proteins where variance primarily reflects biological rather than technical variation, while excluding 553 proteins with S/N < 2 (Figure 2F).

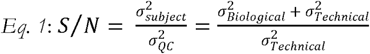

**Figure 2:**
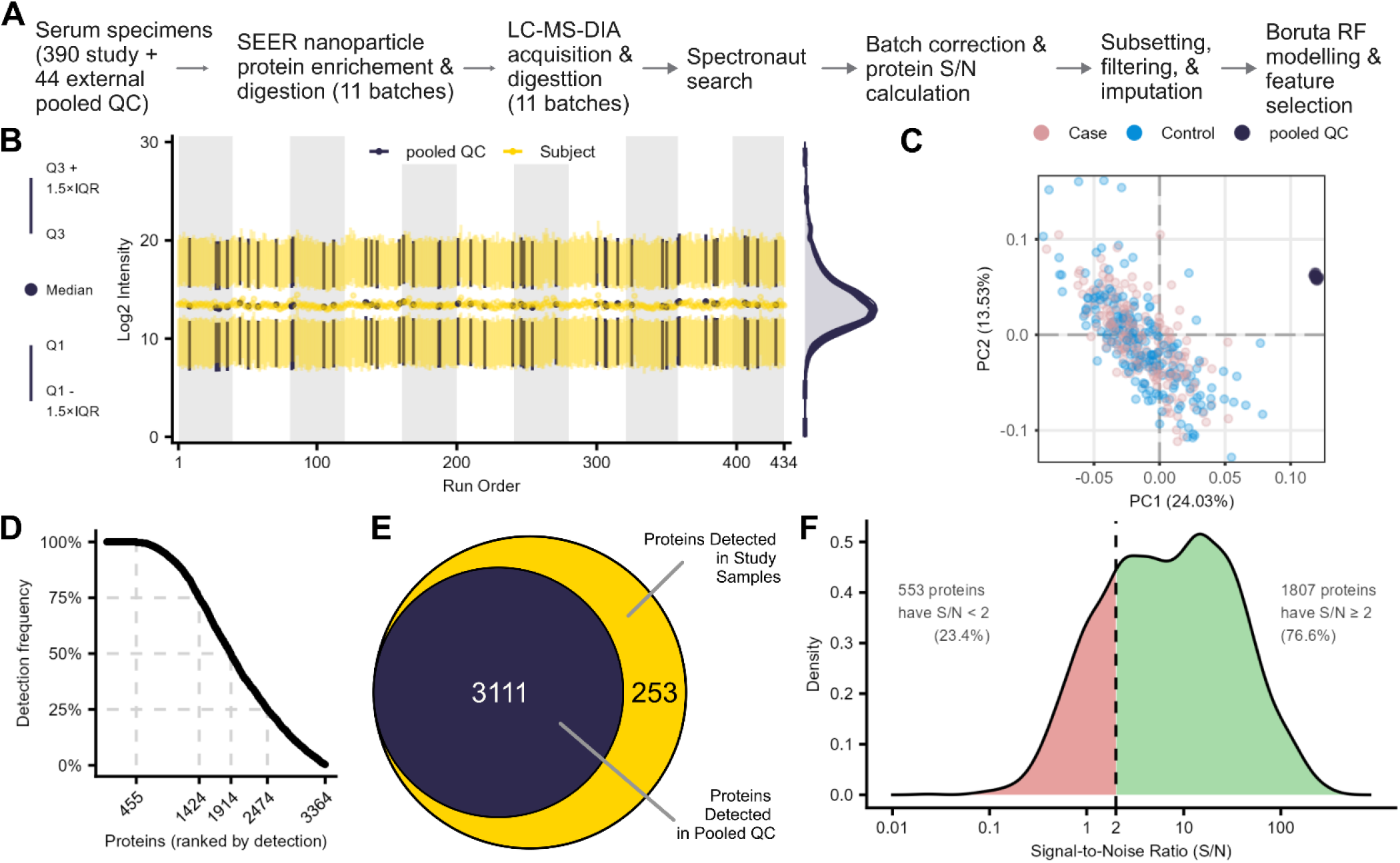
Technical workflow and data quality assessment. (A) Sample processing workflow from serum collection through biomarker discovery. (B) Analytical stability across LC-MS run order; alternating grey and white indicate distinct batches. (C) Principal component analysis of batch-corrected data colored by sample type. (D) Protein detection frequency across samples, showing percentage of proteins detected in varying numbers of samples. (E) Euler diagram of protein detection overlap between pooled QC and study samples. (F) Signal-to-noise ratio distribution for proteins detected in ≥ 25% of pooled QC samples; 76.6% of these have S/N ≥ 2 (dashed line).

Because each analytical comparison involved different subsets of samples, subsequent filtering and preprocessing steps were performed separately for each comparison. This comparison-specific approach was necessary to avoid losing biologically relevant proteins that might be highly present within a specific timepoint or group but rare across the full dataset. Applying global presence filtering would inappropriately exclude proteins that show selective expression patterns in particular biological contexts despite being potentially important markers.

For each analytical comparison, the following preprocessing pipeline was applied independently. For crosssectional analyses comparing cases and controls within the same timepoint, data were first subsetted to include only samples from the relevant timepoint. Proteins were then filtered using the globally calculated signal-to-noise ratios, retaining those with S/N ≥ 2.0. A presence threshold was subsequently applied, requiring detection in at least 50% of samples across both cases and controls combined. Remaining missing values were imputed using k-nearest neighbors (KNN, k=10) [37,38], and data were zero-centered by subtracting each sample’s median log_₂_ protein abundance value.

For longitudinal analyses examining protein abundance changes over time (Δ expression), both timepoints were processed jointly to ensure the same protein panel across measurements. The combined dataset underwent identical S/N (≥ 2.0) and presence (≥ 50%) filtering, followed by KNN imputation. After joint preprocessing, time points were split, and subjects were matched by study identifier (Figure 1C). Only subjects with data at both timepoints were retained for delta calculation, computed as the difference between the later and earlier timepoint (e.g., log_₂_ DX window minus log_₂_ DX_2yr_prior). Delta values were subsequently zerocentered for modeling.

### Biomarker signature discovery using random forest approach

For biomarker discovery, random forest classification was performed using leave-one-out cross-validation (LOOCV) to classify case versus control status. To address the high-dimensional nature of our DIA proteomics data, where the number of features far exceeds sample size (p >> n problem), a two-stage feature selection approach was implemented within each LOOCV fold to prevent data leakage. First, univariate filtering using limma[39] with moderated t-statistics ranked proteins by differential expression, selecting the top 100 features. Clinical cofactors (age, service branch) were appended to the limma-selected protein features when evaluating cofactor models. Second, these prefiltered features (with or without cofactors) underwent multivariate feature selection using Boruta random forest algorithm[18,40–42], which identifies features with predictive importance significantly greater than random shadow features. Final random forest models[43] were trained using Boruta-selected features.

Hyperparameters, including the number of top limma-prefiltered features and the maximum number of Boruta iterations, were optimized through systematic grid search on cross-sectional DX window data, evaluating varying numbers of prefiltered features and maximum Boruta iterations (Figure S7A). Optimal parameters (100 limma features, 100 Boruta runs) were selected based on maximizing area under the receiver operating characteristic curve (AUC). Model stability was assessed across 10 random seeds, yielding an AUC coefficient of variation of 1.1% (Figure S7B). Model performance was assessed using receiver operating characteristic (ROC) analysis,[44] with emphasis placed on AUC values and feature selection consistency across cross-validation folds. Statistical comparisons between models (e.g., protein-only versus protein plus cofactors) were performed using DeLong’s test for correlated ROC curves.[45] AUC 95% confidence intervals (DeLong method) and DeLong tests were computed with the pROC package.[46] The minimum AUC detectable against chance at each sample size (80% power, two-sided α = 0.05) was calculated with the formula of Obuchowski et al.,[47] as implemented in pROC. From LOOCV results, consensus models were built using features selected in ≥50% of cross-validation folds. SHAP (SHapley Additive exPlanations) values[48] were calculated from consensus models to quantify individual feature contributions to predictions.

## Results

### Study Cohort Characteristics

We identified 73 melanoma cases meeting eligibility criteria with sufficient archived serum for analysis, each matched 1:1 to a control (Table 1). Cases and controls were matched by race/ethnicity, age (±1 year), sex, and year of serum sample acquisition. The cohort was predominantly white (90.4%) and male (84.9%), with a median age at diagnosis of 38 years (IQR: 27–45). Among cases, pathologic stage distribution included Stage I (39.7%), Stage II (21.9%), Stage III (30.1%), and Stage IV (2.7%), with 5.5% unknown (Table 2).

**Table 1:** Demographic characteristics of the study cohort. Complete demographic data for 73 matched melanoma case-control pairs (146 total subjects, 390 serum samples) from the DODSR. Age is reported at diagnosis (cases) or corresponding matched timepoint (controls).

| Variable | Case | Control |
|---|---|---|
| Count of subjects | 73 (50.0%) | 73 (50.0%) |
| Age at diagnosis<sup>a</sup> |  |  |
| Mean ± SD | 37.5 ± 11.5 | 37.4 ± 11.3 |
| Median [IQR] | 38 [27-45] | 38 [27-45] |
| Sex |  |  |
| Male | 62 (84.9%) | 62 (84.9%) |
| Female | 11 (15.1%) | 11 (15.1%) |
| Race and ethnicity |  |  |
| Asian/Pacific Islander | 1 (1.4%) | 1 (1.4%) |
| Black | 1 (1.4%) | 1 (1.4%) |
| Hispanic (any race) | 1 (1.4%) | 1 (1.4%) |
| Other | 2 (2.7%) | 2 (2.7%) |
| White | 66 (90.4%) | 66 (90.4%) |
| Unknown | 2 (2.7%) | 2 (2.7%) |
| Service branch<sup>b</sup> |  |  |
| Air Force | 23 (31.5%) | 24 (32.9%) |
| Army | 23 (31.5%) | 26 (35.6%) |
| Coast Guard | 2 (2.7%) | 0 (0.0%) |
| Marine Corps | 2 (2.7%) | 10 (13.7%) |
| Navy | 15 (20.5%) | 13 (17.8%) |
| Unknown | 8 (11.0%) | 0 (0.0%) |
| Samples per timepoint<sup>c</sup> | n = 195 | n = 195 |
| 4 years prior | 53 | 53 |
| 2 years prior | 66 | 66 |
| Diagnosis window | 35 | 35 |
| 2 years post | 41 | 41 |

<sup>a</sup> Age at diagnosis: matched pairs equivalent (paired t-test, p = 0.225)
<sup>b</sup> Service branch was not a matching variable in subject selection
<sup>c</sup> Sample counts: all other rows are subject-level (n = 73 per group)

**Table 2:** Clinicopathologic characteristics of melanoma cases. AJCC pathologic stage collapsed to major categories due to variable registry reporting granularity. Breslow thickness is reported as median [IQR]. Histologic subtype and anatomic site per ICD-O-3 morphology and topography codes.

| Characteristic | Value |
|---|---|
| AJCC pathologic stage |  |
| I (excludes IA) | 29 (39.7%) |
| II (includes IIA, IIB, IIC) | 16 (21.9%) |
| III (includes IIIA, IIIB, IIIC) | 22 (30.1%) |
| IV | 2 (2.7%) |
| Unknown | 4 (5.5%) |
| Median Breslow thickness [IQR] (n=62)□ | 1.71 [1.26–2.30] mm |
| Histologic subtype |  |
| Melanoma, NOS (8720/3) | 41 (56.2%) |
| Superficial spreading (8743/3) | 17 (23.3%) |
| Nodular (8721/3) | 15 (20.5%) |
| Anatomic site |  |
| Trunk (C44.5) | 31 (42.5%) |
| Upper limb (C44.6) | 15 (20.5%) |
| Lower limb (C44.7) | 6 (8.2%) |
| Head/neck (C44.2–C44.4) | 21 (28.8%) |

□ Available for 62/73 cases (85%); 11 cases had missing or unrecorded values in the cancer registry.

Serum samples were requested at four timepoints relative to diagnosis: approximately 4 years prior, 2 years prior, within 1 year after diagnosis (diagnosis window), and 2 years post-diagnosis (Figure 1A-B). Due to military deployment schedules and routine collection intervals, sample availability varied across timepoints (Figure 1C). In total, 390 samples were analyzed. The 2-year prior timepoint had the highest availability (132 samples, 66 pairs), while the diagnosis window had the lowest (70 samples, 35 pairs). Only 17 case-control pairs (23%) had complete samples across all four timepoints. This incomplete sampling precluded traditional repeated-measures longitudinal modeling and necessitated treating each timepoint comparison independently (Figure 1D).

### Deep serum proteome profiling with nanoparticle enrichment and DIA-MS

We employed a semi-automated workflow combining Seer Proteograph nanoparticle enrichment with DIA mass spectrometry to achieve deep proteome coverage across all 390 study samples and 44 external QC samples (Figure 2A). The workflow demonstrated excellent analytical stability across the 11 processing batches, with consistent protein intensities observed for both study samples and the four external pooled QC samples included in each batch (Figure 2B). Principal component analysis following batch correction using Com-Bat effectively normalized technical variation while preserving biological differences between cases and controls (Figure 2C, S5, and S6). The separation of external QC samples from the study cohort reflects their commercial origin and distinct biological composition rather than technical artifacts, as these samples showed high internal consistency throughout the study.

Across the entire study, we quantified 3,364 unique protein groups, with 1,914 proteins (56.9%) detected in at least 50% of samples (Figure 2D). Detection frequencies for all samples, each sample type and each modeling comparison are given in Table S1. Comparison of proteins detected in study samples versus pooled QC revealed 3,111 proteins common to both sets, with 253 proteins uniquely identified in study samples that were excluded from signal-to-noise calculations (Figure 2E). Application of our signal-to-noise ratio metric (see Methods) identified 1,807 proteins with S/N ≥ 2, indicating that variance in these protein measurements primarily reflects biological rather than technical variation (Figure 2F). This subset of high-quality proteins, representing 76.6% of the 2,360 proteins detected in at least 25% of QC samples, formed the basis for subsequent biomarker discovery analyses.

### Random forest modeling identifies candidate protein signatures

Table 3 summarizes the results from random forest modeling. Cross-sectional analysis in the diagnosis window (DX window) achieved the highest classification performance with an AUC of 0.823 (95% CI: 0.723-0.923), analyzing 35 matched case-control pairs from 1,581 filtered proteins (Figure 3A). At the default classification threshold of 0.5, sensitivity was 0.71 (95% CI 0.54–0.85) and specificity was 0.80 (95% CI 0.63–0.92). The model identified 18 protein features across cross-validation iterations, with 8 core features selected in ≥50% of folds (Figure 3B): serine protease 1 (PRSS1), glycogen synthase kinase 3 beta (GSK3B), FAM20C Golgi-associated secretory pathway kinase (FAM20C), carboxylesterase 1 (CES1), C-C motif chemokine lig-and 14 (CCL14), EPH receptor A10 (EPHA10), lectin mannose binding 2 (LMAN2), and inter-alpha-trypsin inhibitor heavy chain 1 (ITIH1). All 8 proteins showed statistically significant differential expression between cases and controls (Figure S8). Peptide-level identification evidence for each candidate marker is summarized in Table S2. Two markers (CCL14 and EPHA10) were quantified from a single peptide. Both passed the signal-to-noise filter applied to every modeled protein (S/N 4.9 and 17.9, respectively; Table S2). SHAP value analysis demonstrated that elevated expression (red points) of PRSS1 and reduced expression (blue points) of FAM20C and CES1 contributed most strongly to case classification, while elevated expression of CCL14 and GSK3B in cases relative to controls also contributed to discrimination (Figure 3C). Model performance decreased at earlier timepoints, with AUCs of 0.609 (95% CI: 0.512-0.706) at 2 years prior and 0.566 (95% CI: 0.455-0.676) at 4 years prior to diagnosis (Figures S9, S10). Post-diagnosis performance yielded an AUC of 0.582 (95% CI: 0.456-0.709), but this result potentially reflects treatment-induced perturbations or, alternatively, disease specificity of the signature as patients who are disease-free post-treatment may lose the signal (Figure S11).

**Figure 3:**
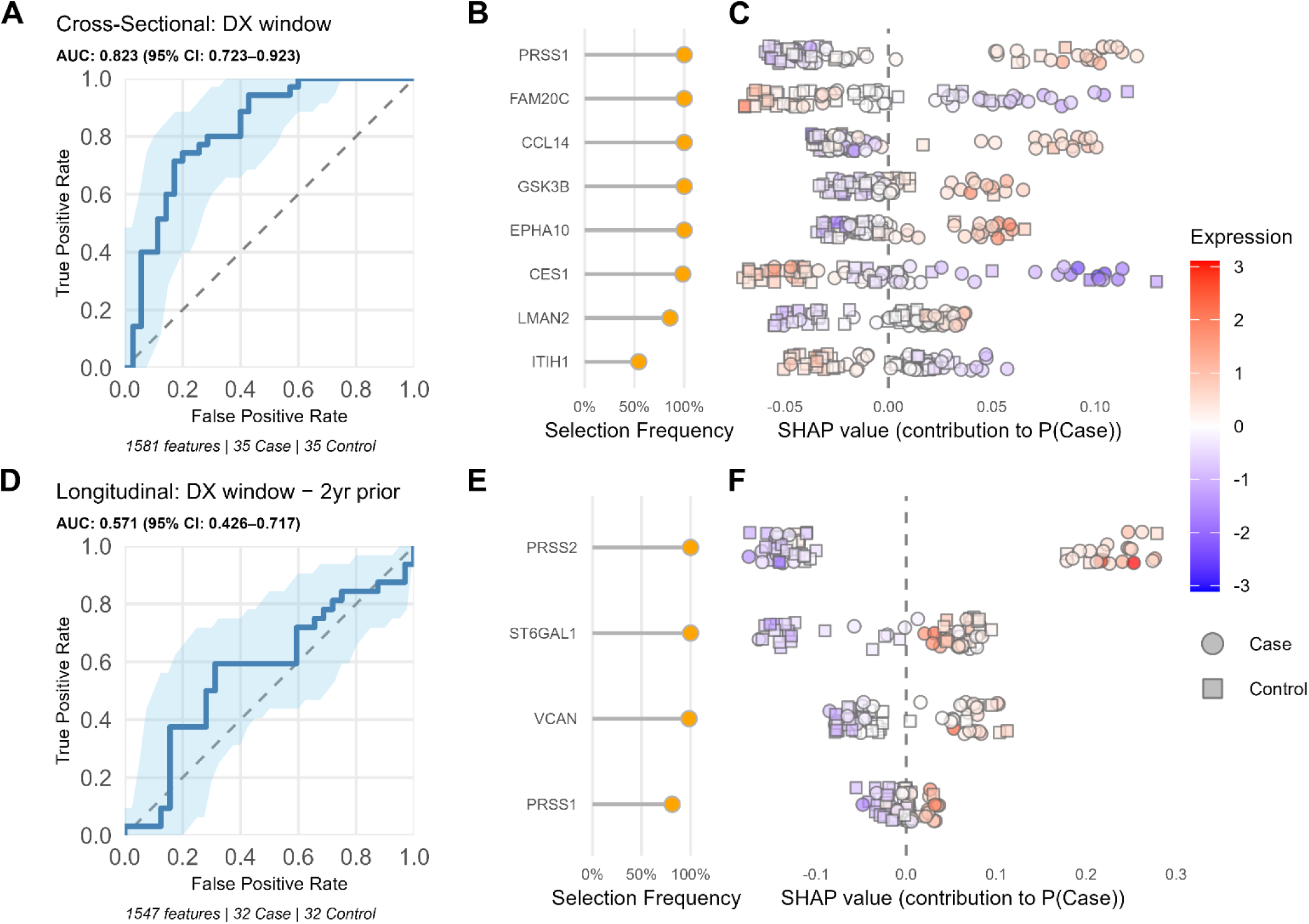
Random forest models for cross-sectional and longitudinal melanoma classification. (A-C) Cross-sectional comparison in the diagnosis window (n=70; 35 cases, 35 controls, 1,581 features). (A) LOOCV ROC curve with 95% CI (AUC = 0.823) from random f rest model with univariate limma prefiltering and Boruta feature selection. (B) Feature selection stability across LOOCV folds. (C) SHAP values from consensus model built using features selected in ≥50% of LOOCV folds, showing contribution to case probability colored by log_2_ ze o-centered protein expression level (red = high, blue = low). Panels B and C share the same y-axis. (D-F) Longitudinal analysis of protein expression changes between the diagnosis window and 2 years prior (n=64; 32 cases, 32 controls, 1,547 features). (D) LOOCV ROC curve (AUC = 0.571). (E) Feature selection stability. (F) SHAP values from consensus model (≥50% selection threshold). Panels E and F share the same y-axis.

**Table 3:**
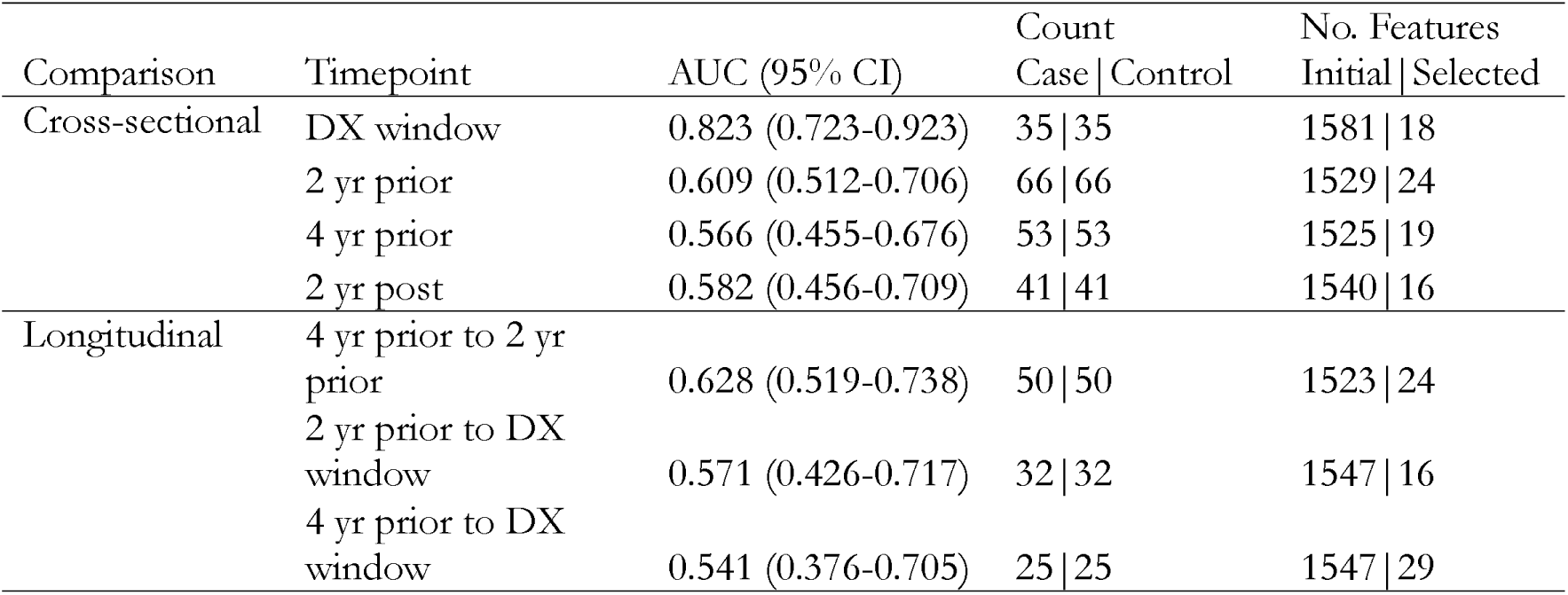
Summary of random forest modeling results. . Note all modeled features are filtered at S/N ≥ 2 and presence ≥ 0.5; AUC is area under the receiver operating characteristic curve. Number of initial features and selected features indicate total number of proteins in that comparison’s input data and features selected in at least one LOOCV fold, respectively.

Longitudinal analysis comparing within-individual protein expression differences between diagnosis and 2 years prior achieved an AUC of 0.571 (95% CI: 0.426-0.717) from 32 matched pairs and 1,547 filtered proteins (Figure 3D). Notably, top consensus features included serine protease 2 (PRSS2) and PRSS1, which were also among the strongest discriminators in the cross-sectional diagnosis-window analysis, alongside ST6 beta-galactoside alpha-2,6-sialyltransferase 1 (ST6GAL1) and versican (VCAN) (Figures 3E, S12). This overlap suggests these proteins both differentiate cases from controls in the diagnosis window and may change over the years preceding diagnosis, although the longitudinal model itself was not distinguishable from chance. SHAP value analysis revealed that increasing PRSS2 and VCAN differences in expression over this interval contributed to case classification (Figure 3F). Other longitudinal comparisons yielded modest discrimination, with the 4-year-prior to 2-year-prior interval achieving an AUC of 0.628 (95% CI: 0.519-0.738) and the 4-year-prior to diagnosis comparison performing near chance (AUC 0.541, 95% CI: 0.376-0.705) (Figures S13, S14).

### Evaluation of demographic variables

To evaluate potential demographic influences on biomarker discovery, we performed sensitivity analyses incorporating age, military service branch, and demographic subset modeling at the DX window timepoint. Rather than including demographic variables as fixed covariates, we appended them to the feature pool postunivariate filtering to test whether they would be selected as discriminative features alongside protein candidates. Adding age as a potential predictor, consistent with our prior DODSR biomarker study[18], yielded an AUC of 0.812 (95% CI: 0.710-0.915, DeLong test p = 0.625, Figure S15), statistically indistinguishable from the 0.823 (95% CI: 0.723–0.923) achieved by proteins alone. Age was never selected as a consensus feature, and the core protein signature (PRSS1, GSK3B, FAM20C, CES1, CCL14, EPHA10, LMAN2, ITIH1) remained unchanged (Figure S15B). Incorporating military service branch likewise did not improve performance (AUC 0.798, 95% CI: 0.691-0.905, DeLong test p = 0.310, Figure S16), and service branch was not selected during feature selection.

To probe generalizability, we re-ran the full discovery pipeline restricted to white males (n = 54), the majority demographic in the cohort. Performance dropped to an AUC of 0.684 (95% CI: 0.541-0.828, Figure S17A), substantially below the 0.823 of the full cohort. Because the two models derive from different samples, their ROC curves cannot be formally compared (precluding a DeLong test), and the reduced sample size and demographic homogeneity of the subset prevent us from distinguishing genuine demographic-specific signal from loss of statistical power. Collectively, these analyses indicate that demographic variables do not improve classification beyond protein features alone; whether the protein signature itself generalizes across demographic groups cannot be resolved in this cohort and remains a question for larger, more diverse validation sets.

## Discussion

We used data-independent acquisition mass spectrometry with nanoparticle enrichment across 390 longitudinal serum samples from melanoma cases and matched controls, quantifying over 3,300 proteins with reproducible measurement. Cross-sectional analysis in the diagnosis window achieved an AUC of 0.823 (95% CI: 0.723-0.923), identifying a consensus signature of 8 proteins (PRSS1, GSK3B, FAM20C, CES1, CCL14, EPHA10, LMAN2, ITIH1), while longitudinal and pre-diagnostic analyses yielded more modest discrimination.

This performance supports further evaluation of serum-based melanoma detection as a complement to clinical examination given the challenges of detecting signals from low tumor mass in complex serum. The consistent selection of serine protease 1 (PRSS1) across both cross-sectional and longitudinal models (the latter featuring PRSS2) is particularly noteworthy, as serine protease activity facilitates tumor invasion through basement membrane degradation and matrix metalloproteinase activation.[49,50] The selection of GSK3B in the cross-sectional consensus signature may further support the signature’s biological relevance, due to GSK3B’s dual roles as tumor suppressor in early stages and invasion promoter in advanced disease[51,52].

The identified candidate markers differ substantially from melanoma tissue proteomics studies, reflecting fundamental differences between tissue and serum analyses.[53] Established advanced-stage serum biomarkers S100B[9] and LDH[10] were detected in our dataset but did not discriminate between cases and controls in the diagnosis window (see Figure S8, LDH p>0.05; S100B did not pass S/N filtering), which is consistent with these markers’ clinical utility being limited to late-stage disease with substantial tumor burden. Our cohort included earlier disease stages where tumor mass is small and direct tumor secretion faces substantial dilution across the total blood volume. Yet we detect proteases, PRSS1 and PRSS2, which could represent tumor-derived signals given serine proteases’ established roles in melanoma invasion and metastasis, although both are also abundant in circulation. This apparent paradox may be resolved by considering these candidates as a mixed panel capturing both tumor-derived factors and systemic host responses. GSK3B plausibly originates from tumor cells and their immediate microenvironment, while PRSS1 and PRSS2, CES1 (carboxylesterase involved in metabolic processing), CCL14 (chemokine potentially indicating immune activation), and ITIH1 (extracellular matrix stabilizer) may reflect systemic host responses to developing malignancy. The secretory-pathway kinase FAM20C and the sialyltransferase ST6GAL1, a longitudinal feature, suggest post-translational modification pathways generate detectable systemic signatures even when absolute tumorderived protein abundance remains low. This interpretation aligns with accumulating evidence that early cancer detection in biofluids often reflects host response rather than direct tumor secretion, particularly for solid tumors with limited vascular access.[54] Because our controls were free of cancer, we cannot determine whether this signature is specific to melanoma; several candidates, including PRSS1, CES1 and GSK3B, have been associated with other cancers,[51,52,55–57] and comparison with other malignancies and with benign skin lesions will be needed to establish specificity.

Model performance declined progressively at earlier timepoints (AUC 0.609 at 2 years prior, 0.566 at 4 years prior) suggesting discriminative protein changes occur primarily proximal to clinical diagnosis. Longitudinal analysis comparing within-individual changes from 2 years prior to diagnosis achieved more modest performance (AUC 0.571, 95% CI: 0.426-0.717). However, PRSS1 and PRSS2 recurred across cross-sectional and longitudinal models, although the two analyses share the diagnosis-window samples. The modest longitudinal performance contrasts with more dramatic longitudinal improvements in our group’s previous oropharyngeal squamous cell carcinoma study[18] (AUC 0.90 longitudinal vs. 0.76 cross-sectional), possibly reflecting the early cancer stage of our melanoma cohort and suggesting melanoma’s systemic protein signatures may evolve on a shorter timespan than our previous studies’ 2-year sample collection window.

Sensitivity analyses incorporating age and military service branch provided no improvement over proteinonly models (age: AUC 0.812, 95% CI: 0.710-0.915, p = 0.625; service branch: AUC 0.798, 95% CI: 0.6910.905, p = 0.310 vs. proteins: AUC 0.823), and neither variable was selected during feature selection despite being available, indicating the protein signal does not simply encode these demographic factors. Generalizability across demographic groups is less settled: restricting the analysis to white males reduced performance to an AUC of 0.684 (95% CI: 0.541-0.828), but the smaller, more homogeneous subset (n = 54) cannot separate genuine demographic-specific effects from reduced statistical power. Sample size likewise precluded meaningful stratification by disease characteristics such as tumor stage or anatomic site. Establishing whether the signature holds across race and sex will require larger, more balanced validation cohorts.

Our study population, predominantly white, male, military personnel (84.9% male), presents limitations for generalizability. This composition reflects well-documented patterns in melanoma epidemiology and military demographics. Melanoma incidence is substantially higher in white populations,[1] and military cohorts experience unique environmental exposures including intense UV radiation and insufficient sun protection during deployment.[58–60] Female underrepresentation (15.1% of cases), reflecting underrepresentation of females in the active-duty population (17.7% in 2023[61]), limits identification of sex-specific biomarkers.

These characteristics emphasize the need for future validation in larger, more diverse cohorts with balanced representation across demographic groups and disease stages. Additional limitations include incomplete sampling across timepoints driven by military deployment schedules and variable duration of active-duty service, which prevented traditional longitudinal modeling and may have obscured complex temporal patterns. The cohort size was set by the availability of archived specimens rather than by an a priori power calculation. With 35 matched pairs, the smallest true AUC the study could distinguish from chance with 80% power is approximately 0.69 (0.64 with the 66 pairs available 2 years before diagnosis), so the weak pre-diagnostic results should be read as inconclusive rather than as evidence that no pre-diagnostic signal exists.

Our analysis strategy prioritized feature robustness over sensitivity, potentially excluding biologically relevant but technically variable proteins. We lacked patient-level risk factor data, including sun exposure history, sunscreen use, and deployment locations, that would enable correlation of candidate markers with environmental exposures. We also lacked treatment details for the post-diagnosis timepoint, limiting interpretation of the near-chance performance (AUC 0.582) at that interval. All samples were serum; per-specimen DODSR collection records were not retrieved for this study. Erythrocyte, platelet and coagulation marker panels did not differ between cases and matched controls at any timepoint (Figure S18), and none of the candidate markers is among proteins in these marker panels or the proteins that differ most between plasma and serum.[62]

Despite these limitations, the study has notable strengths: pre-diagnostic serum from the DODSR, which is rarely available for melanoma; a matched case-control design; and deep proteome coverage. Our workflow incorporated external pooled quality control samples to calculate signal-to-noise ratios for each protein, distinguishing biological variance from technical variance and reducing the likelihood that selected proteins reflect analytical artifacts. Even if these function primarily as diagnostic markers rather than early detection markers, they could benefit patients with suspicious lesions who wish to avoid invasive biopsy, such as when lesions are present in sensitive areas like the face, and patients with diagnostically challenging lesions, although this use would need to be tested in samples collected before biopsy. A positive result using our panel would prompt careful full-body skin examination, sampling of atypical lesions, and increased surveillance, rather than immediate definitive diagnosis. Clinical trials with sufficient follow-up will be required to determine the clinical utility of this panel.

Validation in larger, independent cohorts is essential to establish reproducibility and assess performance across diverse demographic groups and disease stages. Future studies should incorporate comprehensive clinical annotation, tumor characteristics, and recurrence outcomes to determine whether the candidate markers predict disease severity or clinical trajectories beyond initial diagnosis. Such work could inform development of serological screening tools for high-risk populations who might benefit from regular dermatological monitoring, complementing existing surveillance strategies for melanoma detection.

## Conclusions

We identified serum protein signatures distinguishing melanoma cases from controls in the diagnosis window (AUC 0.823) with weak pre-diagnostic and longitudinal signals. While performance does not yet support clinical implementation, these results provide preliminary evidence for serum-based melanoma detection and identify candidate markers for validation in independent patient cohorts. The consistent selection of serine proteases (PRSS1, PRSS2) alongside markers of post-translational modification, immune activation, and extracellular matrix remodeling suggests these biological processes generate detectable systemic signatures at or near diagnosis, potentially reflecting combined tumor-derived factors and host responses to developing malignancy.

## List of abbreviations

ACTUR: Automated Central Tumor Registry
AJCC: American Joint Committee on Cancer
AUC: Area under the receiver operating characteristic curve
CCL14: C-C motif chemokine ligand 14
CES1: Carboxylesterase 1
CI: Confidence interval
ctDNA: Circulating tumor DNA
DIA: Data-independent acquisition
DoD: Department of Defense
DODSR: Department of Defense Serum Repository
DX: Diagnosis; used in sample timepoint labels
EPHA10: EPH receptor A10
FAM20C: FAM20C Golgi-associated secretory pathway kinase
GSK3B: Glycogen synthase kinase 3 beta
ICD-O-3: International Classification of Diseases for Oncology, Third Edition
iRT: Indexed retention time; synthetic peptide standards spiked into samples to normalize chromatographic retention time
ITIH1: Inter-alpha-trypsin inhibitor heavy chain 1
JPC: Joint Pathology Center
KNN: k-nearest neighbors
LC-MS: Liquid chromatography-mass spectrometry
LDH: Lactate dehydrogenase
LMAN2: Lectin mannose binding 2
LOOCV: Leave-one-out cross-validation
MCED: Multi-cancer early detection
NMSC: Non-melanoma skin cancer
NP-A / NP-B: The two engineered nanoparticle pools of the Seer Proteograph XT assay
PCA: Principal component analysis
PRSS1: Serine protease 1
PRSS2: Serine protease 2
QC: Quality control
ROC: Receiver operating characteristic
S100B: S100 calcium-binding protein B
S/N: Signal-to-noise ratio, calculated here as the ratio of variance across study samples to variance across pooled quality control samples
SHAP: SHapley Additive exPlanations; method for quantifying individual feature contributions to model predictions
SPD: Samples per day; throughput setting of the liquid chromatography gradient method
ST6GAL1: ST6 beta-galactoside alpha-2,6-sialyltransferase 1
USUHS: Uniformed Services University of the Health Sciences VCAN: Versican

## Declarations

## Ethics approval and consent to participate

This study was approved by the Uniformed Services University of the Health Sciences Human Research Protections Program (Protocol No. DBS.2020.119) and determined to be research not involving human subjects as defined at 32 CFR 219.102(e)(1) and applicable DoD policy guidance. Accordingly, informed consent was not required.

## Consent for publication

Not applicable.

## Availability of data and materials

The mass spectrometry proteomics data generated in this study, comprising 434 DIA-MS runs (390 study samples and 44 pooled QC samples), and associated metadata and search results, have been deposited in the MassIVE repository[63] (MSV000100913) and are publicly available.

## Authors’ contributions

C.D.S., K.D.R. and T.L. conceived and designed the study. J.C.A.L. coordinated sample acquisition and cohort assembly, including DODSR access and registry and clinical data. D.R.H., B.I.G., and A.A.S. performed serum sample preparation and proteomics processing. D.R.H. and T.L.F. performed LC-MS data acquisition. D.R.H. carried out the computational, machine-learning, and statistical analyses, incorporating code and advice from B.D. and V.A.P., with minor code contributions from T.J.S. J.C.A.L., L.S.G., E.S.K. and I.B. led clinical interpretation, with additional input from C.L.D. and J.S.H.L. D.R.H. wrote the original draft. T.L. and V.A.P. edited the manuscript, with input from all co-authors. C.D.S. and T.L. acquired funding and supervised the study. All authors reviewed and approved the final manuscript.

## Acknowledgements

The authors thank the clinical and laboratory staff at USUHS and PNNL.

The contents of this publication are the sole responsibility of the author(s) and do not necessarily reflect the views, opinions or policies of USUHS, The Henry M. Jackson Foundation for the Advancement of Military Medicine, Inc., the Department of War, the Departments of the Army, Navy, or Air Force. Mention of trade names, commercial products, or organizations does not imply endorsement by the U.S. Government.

The contributions of the NIH author are considered Works of the United States Government. The findings and conclusions presented in this paper are those of the authors and do not necessarily reflect the views of the NIH or the U.S. Department of Health and Human Services.

## Supplementary Information

See the supplementary information for details on study design and matching criteria. Supplementary Data contains sample metadata (sheet “Sample Metadata”), raw protein group-level expression values from Spectronaut processing (sheet “Raw Expression”), and batch-corrected protein group-level expression values used in downstream analyses (sheet “Batch Corrected Expression”), provided as a multi-sheet Excel workbook.

## References

1. Siegel RL, Kratzer TB, Wagle NS, Sung H, Jemal A. Cancer statistics, 2026. CA Cancer J Clin. 2026;76:e70043. 10.3322/caac.70043

2. Lee T, Williams VF, Clark LL. Incident diagnoses of cancers in the active component and cancer-related deaths in the active and reserve components, U.S. Armed Forces, 2005-2014. MSMR. 2016;23:23–31.

3. Merlino G, Herlyn M, Fisher DE, Bastian BC, Flaherty KT, Davies MA, et al. The state of melanoma: challenges and opportunities. Pigment Cell Melanoma Res. 2016;29:404–16. 10.1111/pcmr.12475

4. Zaremba A, Lodde G, Murali R, Philip M, Cosgarea I, Jansen P, et al. Molecular pathology as a diagnostic aid in difficult to classify melanocytic tumors with spitzoid morphology. Eur J Cancer Oxf Engl 1990. 2021;148:340–7. 10.1016/j.ejca.2021.02.025

5. National Cancer Institute Clinical Proteomic Tumor Analysis Consortium (CPTAC). The Clinical Proteomic Tumor Analysis Consortium Cutaneous Melanoma Collection (CPTAC-CM) [Internet]. The Cancer Imaging Archive; 2018 [cited 2025 Oct 8]. 10.7937/K9/TCIA.2018.ODU24GZE

6. Guedes J, Szadai L, Woldmar N, Jánosi ÁJ, Koroncziová K, Lengyel BM, et al. The melanoma MEGAstudy: Integrating proteogenomics, digital pathology, and AI-analytics for precision oncology. J Proteomics. 2025;319:105482. 10.1016/j.jprot.2025.105482

7. Betancourt LH, Gil J, Kim Y, Doma V, Çakır U, Sanchez A, et al. The human melanoma proteome atlas-Defining the molecular pathology. Clin Transl Med. 2021;11:e473. 10.1002/ctm2.473

8. Pekarek L, Cedra AS, Jaudenes YDY, Ospino LR, Pedrejón BI, Bernier L, et al. Paradigm of biomarkers in metastatic melanoma (Review). Oncol Lett. Spandidos Publications; 2025;29:1–12. 10.3892/ol.2024.14824

9. Gaynor R, Irie R, Morton D, Herschman HR, Jones P, Cochran A. S100 PROTEIN: A MARKER FOR HUMAN MALIGNANT MELANOMAS? The Lancet. 1981;317:869–71. 10.1016/S0140-6736(81)92142-5

10. Xu J, Zhao J, Wang J, Sun C, Zhu X. Prognostic value of lactate dehydrogenase for melanoma patients receiving anti-PD-1/PD-L1 therapy. Medicine (Baltimore). 2021;100:e25318. 10.1097/MD.0000000000025318

11. Bartolomucci A, Nobrega M, Ferrier T, Dickinson K, Kaorey N, Nadeau A, et al. Circulating tumor DNA to monitor treatment response in solid tumors and advance precision oncology. Npj Precis Oncol. Nature Publishing Group; 2025;9:84. 10.1038/s41698-025-00876-y

12. Imai M, Nakamura Y, Yoshino T. Transforming cancer screening: the potential of multi-cancer early detection (MCED) technologies. Int J Clin Oncol. 2025;30:180–93. 10.1007/s10147-025-02694-5

13. Guzman UH, Martinez-Val A, Ye Z, Damoc E, Arrey TN, Pashkova A, et al. Ultra-fast label-free quantification and comprehensive proteome coverage with narrow-window data-independent acquisition. Nat Biotechnol. Nature Publishing Group; 2024;42:1855–66. 10.1038/s41587-023-02099-7

14. Fröhlich K, Fahrner M, Brombacher E, Seredynska A, Maldacker M, Kreutz C, et al. Data-Independent Acquisition: A Milestone and Prospect in Clinical Mass Spectrometry-Based Proteomics. Mol Cell Proteomics MCP. 2024;23:100800. 10.1016/j.mcpro.2024.100800

15. Blume JE, Manning WC, Troiano G, Hornburg D, Figa M, Hesterberg L, et al. Rapid, deep and precise profiling of the plasma proteome with multi-nanoparticle protein corona. Nat Commun. Nature Publishing Group; 2020;11:3662. 10.1038/s41467-020-17033-7

16. Huang T, Wang J, Stukalov A, Kreimer S, Zhao X, Cantrell LS, et al. Multiplexed Nanoparticle Protein Corona Enables Accurate and Precise Deep Plasma Proteomics. J Proteome Res. 2025;24:5884–93. 10.1021/acs.jproteome.5c00729

17. Perdue CL, Cost AAE, Rubertone MV, Lindler LE, Ludwig SL. Description and Utilization of the United States Department of Defense Serum Repository: A Review of Published Studies, 1985-2012. PLOS ONE. Public Library of Science; 2015;10:e0114857. 10.1371/journal.pone.0114857

18. Lee JY, Shi T, Petyuk VA, Schepmoes AA, Fillmore TL, Wang Y-T, et al. Detection of Head and Neck Cancer Based on Longitudinal Changes in Serum Protein Abundance. Cancer Epidemiol Biomarkers Prev. 2020;29:1665–72. 10.1158/1055-9965.EPI-20-0192

19. Pease PJ, Jiang W, Zhao X, Benz R, Williamson L, Chu J, et al. Evaluation of blood-based sample types for deep plasma proteomics [Internet]. Redwood City, CA: Seer Inc.; 2023 Feb. Report No.: CF-1049 REV A. https://media.seer.bio/uploads/2023/02/serum-v-plasma-appnote.pdf

20. Suhre K, Chen Q, Halama A, Mendez K, Dahlin A, Stephan N, et al. A genome-wide association study of mass spectrometry proteomics using a nanoparticle enrichment platform. Nat Genet. Nature Publishing Group; 2025;57:2987–96. 10.1038/s41588-025-02413-w

21. Ferdosi S, Tangeysh B, Brown TR, Everley PA, Figa M, McLean M, et al. Engineered nanoparticles enable deep proteomics studies at scale by leveraging tunable nano–bio interactions. Proc Natl Acad Sci. 2022;119:e2106053119. 10.1073/pnas.2106053119

22. Chen S-Y, Williamson L, Donovan MKR, Gajadhar AS. Balancing deep proteome coverage with limited sample amounts using Seer’s Proteograph Product Suite [Internet]. Seer Inc.; 2023. Report No.: CF-1050 REV A. https://media.seer.bio/uploads/2023/02/dilution-low-volume-appnote.pdf. Accessed 10 Oct 2025

23. Roger K, Metatla I, Ceccacci S, Wahbi K, Motté L, Chhuon C, et al. Mining the plasma proteome: Evaluation of enrichment methods for depth and reproducibility. J Proteomics. 2025;321:105519. 10.1016/j.jprot.2025.105519

24. Kverneland AH, Harking F, Vej-Nielsen JM, Huusfeldt M, Bekker-Jensen DB, Svane IM, et al. Fully Automated Workflow for Integrated Sample Digestion and Evotip Loading Enabling High-Throughput Clinical Proteomics. Mol Cell Proteomics [Internet]. Elsevier; 2024 [cited 2025 May 7];23. 10.1016/j.mcpro.2024.100790

25. Jumel T, Shevchenko A. Multispecies Benchmark Analysis for LC-MS/MS Validation and Performance Evaluation in Bottom-Up Proteomics. J Proteome Res. American Chemical Society; 2024;23:684–91. 10.1021/acs.jproteome.3c00531

26. Pino LK, Searle BC, Yang H-Y, Hoofnagle AN, Noble WS, MacCoss MJ. Matrix-Matched Calibration Curves for Assessing Analytical Figures of Merit in Quantitative Proteomics. J Proteome Res. 2020;19:1147–53. 10.1021/acs.jproteome.9b00666

27. Bache N, Geyer PE, Bekker-Jensen DB, Hoerning O, Falkenby L, Treit PV, et al. A Novel LC System Embeds Analytes in Pre-formed Gradients for Rapid, Ultra-robust Proteomics*. Mol Cell Proteomics. 2018;17:2284–96. 10.1074/mcp.TIR118.000853

28. Westgard JO. Basic QC practices: training in statistical quality control for medical laboratories. Fourth edition. Madison, WI: Westgard QC; 2016.

29. Bruderer R, Bernhardt OM, Gandhi T, Miladinović SM, Cheng L-Y, Messner S, et al. Extending the Limits of Quantitative Proteome Profiling with Data-Independent Acquisition and Application to Acetaminophen-Treated Three-Dimensional Liver Microtissues*[S]. Mol Cell Proteomics. 2015;14:1400–10. 10.1074/mcp.M114.044305

30. Mudge JM, Carbonell-Sala S, Diekhans M, Martinez JG, Hunt T, Jungreis I, et al. GENCODE 2025: reference gene annotation for human and mouse. Nucleic Acids Res. 2025;53:D966–75. 10.1093/nar/gkae1078

31. Gupta N, Pevzner PA. False Discovery Rates of Protein Identifications: A Strike against the Two-Peptide Rule. J Proteome Res. 2009;8:4173–81. 10.1021/pr9004794

32. UniProt Consortium. UniProt: the universal protein knowledgebase in 2021. Nucleic Acids Res. 2021;49:D480–9. 10.1093/nar/gkaa1100

33. Smedley D, Haider S, Ballester B, Holland R, London D, Thorisson G, et al. BioMart – biological queries made easy. BMC Genomics. 2009;10:22. 10.1186/1471-2164-10-22

34. Callister SJ, Barry RC, Adkins JN, Johnson ET, Qian W, Webb-Robertson B-JM, et al. Normalization Approaches for Removing Systematic Biases Associated with Mass Spectrometry and Label-Free Proteomics. J Proteome Res. American Chemical Society; 2006;5:277–86. 10.1021/pr050300l

35. Han W, Hu C, Fan Z-J, Shen G-L. Transcript levels of keratin 1/5/6/14/15/16/17 as potential prognostic indicators in melanoma patients. Sci Rep. Nature Publishing Group; 2021;11:1023. 10.1038/s41598-020-80336-8

36. Johnson WE, Li C, Rabinovic A. Adjusting batch effects in microarray expression data using empirical Bayes methods. Biostatistics. 2007;8:118–27. 10.1093/biostatistics/kxj037

37. Hastie T, Tibshirani R. impute [Internet]. Bioconductor; 2017 [cited 2025 Oct 10]. 10.18129/B9.BIOC.IMPUTE

38. Wei R, Wang J, Su M, Jia E, Chen S, Chen T, et al. Missing Value Imputation Approach for Mass Spectrometry-based Metabolomics Data. Sci Rep. Nature Publishing Group; 2018;8:663. 10.1038/s41598-017-19120-0

39. Ritchie ME, Phipson B, Wu D, Hu Y, Law CW, Shi W, et al. limma powers differential expression analyses for RNA-sequencing and microarray studies. Nucleic Acids Res. 2015;43:e47. 10.1093/nar/gkv007

40. Kursa MB, Rudnicki WR. Feature Selection with the **Boruta** Package. J Stat Softw [Internet]. 2010 [cited 2025 Oct 10];36. 10.18637/jss.v036.i11

41. Breiman L, Cutler A, Liaw A, Wiener M. randomForest: Breiman and Cutlers Random Forests for Classification and Regression [Internet]. 2002 [cited 2025 Oct 10]. p. 4.7-1.2. https://CRAN.R-project.org/package=randomForest. Accessed 10 Oct 2025

42. Breiman L. Random Forests. Mach Learn. 2001;45:5–32. 10.1023/A:1010933404324

43. Wright MN, Ziegler A. ranger: A Fast Implementation of Random Forests for High Dimensional Data in C++ and R. J Stat Softw [Internet]. 2017 [cited 2025 Nov 11];77. 10.18637/jss.v077.i01

44. Sing T, Sander O, Beerenwinkel N, Lengauer T. ROCR: visualizing classifier performance in R. Bioinformatics. 2005;21:3940–1. 10.1093/bioinformatics/bti623

45. DeLong ER, DeLong DM, Clarke-Pearson DL. Comparing the areas under two or more correlated receiver operating characteristic curves: a nonparametric approach. Biometrics. 1988;44:837–45.

46. Robin X, Turck N, Hainard A, Tiberti N, Lisacek F, Sanchez J-C, et al. pROC: an open-source package for R and S+ to analyze and compare ROC curves. BMC Bioinformatics. 2011;12:77. 10.1186/1471-2105-12-77

47. Obuchowski NA, Lieber ML, Wians FH. ROC Curves in Clinical Chemistry: Uses, Misuses, and Possible Solutions. Clin Chem. 2004;50:1118–25. 10.1373/clinchem.2004.031823

48. Komisarczyk K, Kozminski P, Maksymiuk S, Biecek P. treeshap: Compute SHAP Values for Your TreeBased Models Using the “TreeSHAP” Algorithm [Internet]. 2023 [cited 2025 Nov 11]. p. 0.3.1. 10.32614/CRAN.package.treeshap

49. Sui L, Wang S, Ganguly D, El Rayes TP, Askeland C, Børretzen A, et al. PRSS2 remodels the tumor microenvironment via repression of Tsp1 to stimulate tumor growth and progression. Nat Commun. 2022;13:7959. 10.1038/s41467-022-35649-9

50. Blackburn JS, Liu I, Coon CI, Brinckerhoff CE. A Matrix Metalloproteinase-1/Protease Activated Receptor-1 signaling axis promotes melanoma invasion and metastasis. Oncogene. 2009;28:4237–48. 10.1038/onc.2009.272

51. Domoto T, Pyko IV, Furuta T, Miyashita K, Uehara M, Shimasaki T, et al. Glycogen synthase kinase-3β is a pivotal mediator of cancer invasion and resistance to therapy. Cancer Sci. 2016;107:1363–72. 10.1111/cas.13028

52. Kao S-H, Wang W-L, Chen C-Y, Chang Y-L, Wu Y-Y, Wang Y-T, et al. GSK3β controls epithelial– mesenchymal transition and tumor metastasis by CHIP-mediated degradation of Slug. Oncogene. Nature Publishing Group; 2014;33:3172–82. 10.1038/onc.2013.279

53. Wu C-C, Hsu C-W, Chen C-D, Yu C-J, Chang K-P, Tai D-I, et al. Candidate Serological Biomarkers for Cancer Identified from the Secretomes of 23 Cancer Cell Lines and the Human Protein Atlas*. Mol Cell Proteomics. 2010;9:1100–17. 10.1074/mcp.M900398-MCP200

54. Connal S, Cameron JM, Sala A, Brennan PM, Palmer DS, Palmer JD, et al. Liquid biopsies: the future of cancer early detection. J Transl Med. 2023;21:118. 10.1186/s12967-023-03960-8

55. Na K, Lee E-Y, Lee H-J, Kim K-Y, Lee H, Jeong S-K, et al. Human plasma carboxylesterase 1, a novel serologic biomarker candidate for hepatocellular carcinoma. Proteomics. 2009;9:3989–99. 10.1002/pmic.200900105

56. Liu Q, Guo L, Zhang S, Wang J, Lin X, Gao F. PRSS1 mutation: a possible pathomechanism of pancreatic carcinogenesis and pancreatic cancer. Mol Med. 2019;25:44. 10.1186/s10020-019-0111-4

57. Yi Q, Dong F, Lin L, Liu Q, Chen S, Gao F, et al. PRSS1 mutations and the proteinase/antiproteinase imbalance in the pathogenesis of pancreatic cancer. Tumor Biol. 2016;37:5805–10. 10.1007/s13277-015-3982-1

58. Armed Forces Health Surveillance Branch. Incident Diagnoses of Malignant Melanoma, Active Components, U.S. Armed Forces, January 1998-June 2008. MSMR. 2008;15:6–9.

59. Wolf ST, Kenney LE, Kenney WL. Ultraviolet Radiation Exposure, Risk, and Protection in Military and Outdoor Athletes. Curr Sports Med Rep. 2020;19:137–41. 10.1249/JSR.0000000000000702

60. Riemenschneider K, Liu J, Powers JG. Skin cancer in the military: A systematic review of melanoma and nonmelanoma skin cancer incidence, prevention, and screening among active duty and veteran personnel. J Am Acad Dermatol. 2018;78:1185–92. 10.1016/j.jaad.2017.11.062

61. Department of Defense. 2023 demographics: Profile of the military community [Internet]. Office of the Deputy Assistant Secretary of Defense for Military Community and Family Policy; 2023. https://download.militaryonesource.mil/12038/MOS/Reports/2023-demographics-report.pdf

62. Geyer PE, Voytik E, Treit PV, Doll S, Kleinhempel A, Niu L, et al. Plasma Proteome Profiling to detect and avoid sample-related biases in biomarker studies. EMBO Mol Med. 2019;11:e10427. 10.15252/emmm.201910427

63. Wang M, Wang J, Carver J, Pullman BS, Cha SW, Bandeira N. Assembling the Community-Scale Discoverable Human Proteome. Cell Syst. 2018;7:412–421.e5. 10.1016/j.cels.2018.08.004

