## Supplementary Information for "Detection of Melanoma Using Deep Serum Proteome Profiling and Machine Learning"

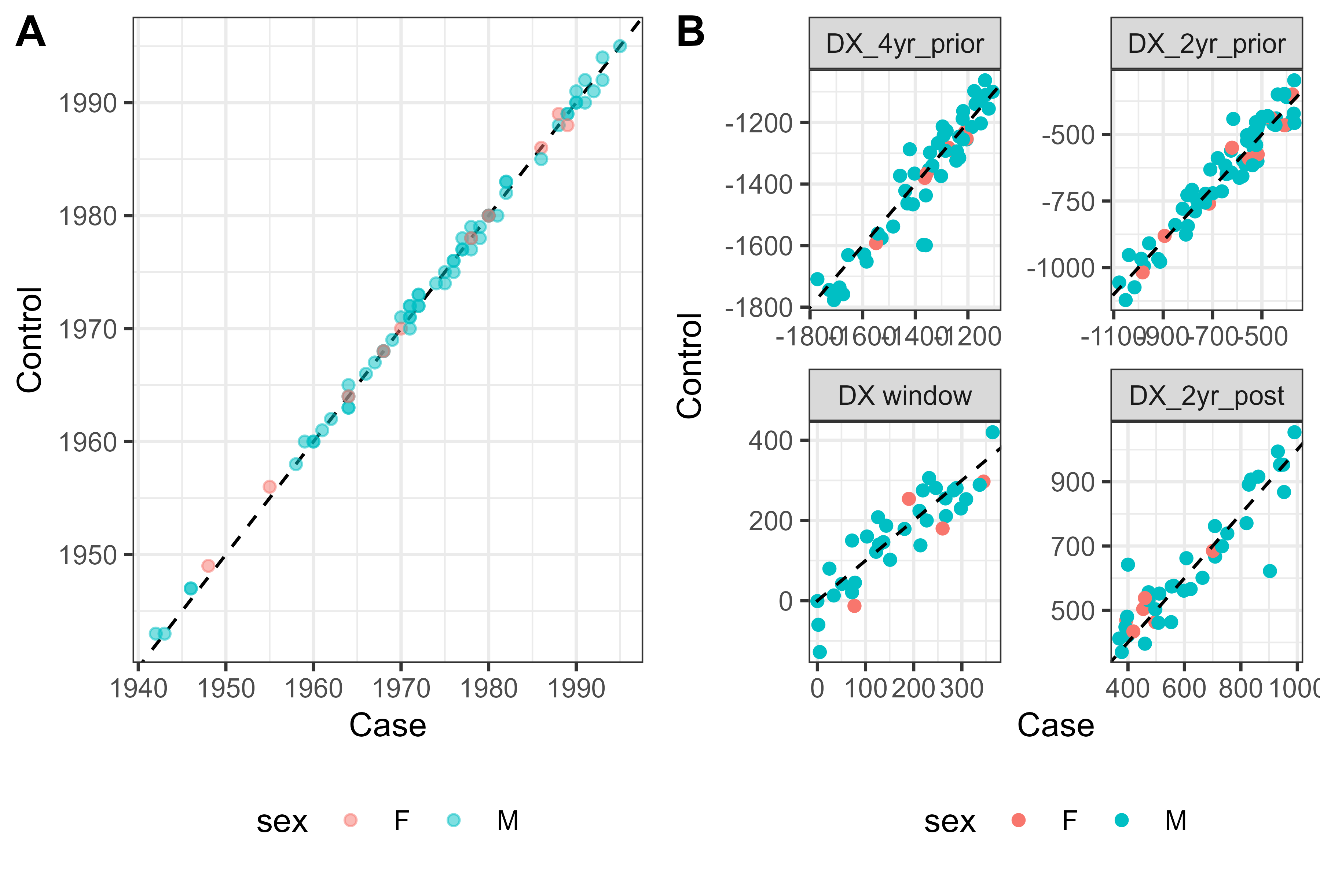

Figure S1: Case-control matching demonstrates tight temporal pairing of samples. (A) Comparison of year of birth between matched case-control pairs, showing excellent age matching (points colored by sex). (B) Comparison of sample collection timing (in days) relative to diagnosis for each timepoint category. Each point represents a matched case-control pair, with x-axis showing the case's collection time and y-axis showing the matched control's collection time relative to their respective reference dates. Dashed lines indicate perfect temporal matching (unity). Close alignment along the unity line demonstrates that case and control samples were collected at comparable time intervals relative to diagnosis across all timepoints (2 years post-diagnosis, 2 years prior, 4 years prior, and diagnosis window).

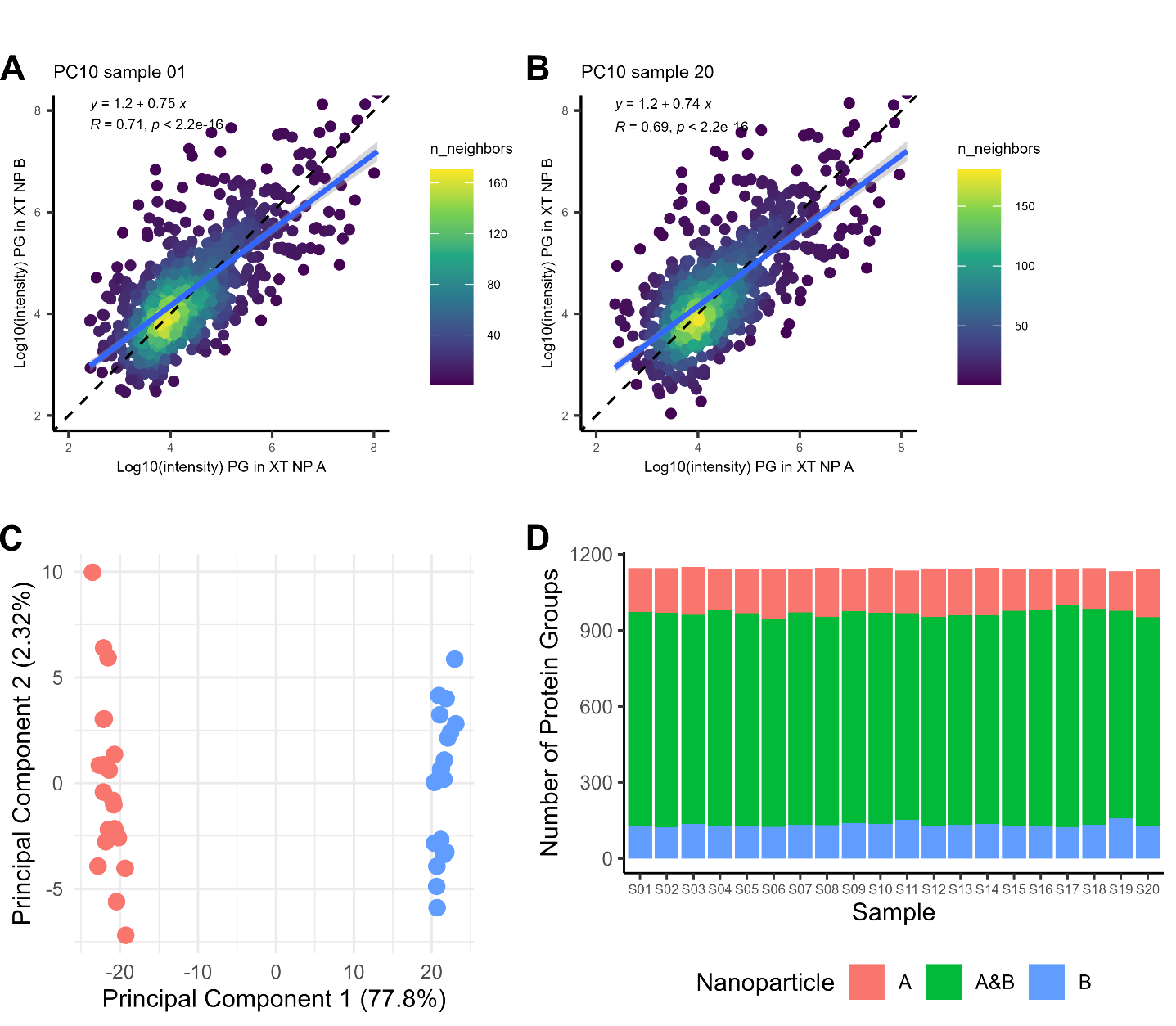

Figure S2: Comparison of protein enrichment between Seer Proteograph XT nanoparticles A and B in n=20 technical replicates of pooled plasma (n = 10 donors, PC10). (A-B) Representative comparison of protein group intensities between NP A and NP B for PC10 sample 01 and sample 20, showing strong correlation (R² = 0.71 and 0.69, respectively; p < 2.2e-16), but systematic intensity differences depending on nanoparticle. Points are colored by local density (n_neighbors) and dashed line indicates unity. (C) PCA plot demonstrates clear separation between NP A (red) and NP B (blue) processed samples across all 20 replicates (40 LC-MS runs total), with PC1 capturing 77.8% of variance. (D) Stacked bar chart showing the number of protein groups detected in each sample, categorized by nanoparticle type: NP A only (red), both A&B (green), and NP B only (blue). PC10 samples were processed using the Seer Proteograph XT assay with 500 ng on column from either NP A or NP B peptides and analyzed on an Evosep One system (60 SPD) with DIA acquisition on an Exploris 480 MS.

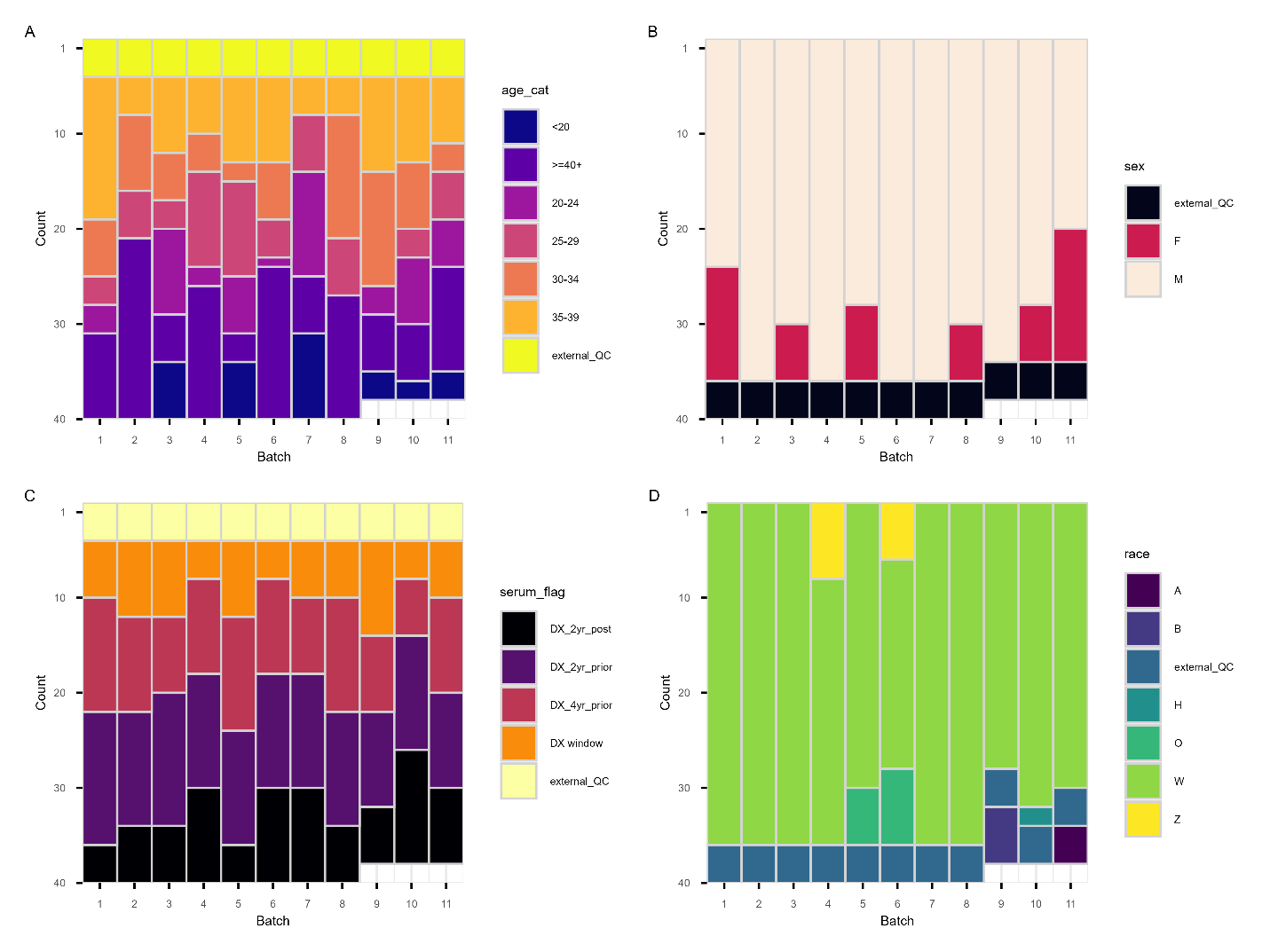

Figure S3: Distribution of sample characteristics across Seer processing batches demonstrates successful randomization. Stacked bar charts showing the composition of each of the 11 Seer batches by (A) age category, (B) sex, (C) sample timepoint relative to diagnosis, and (D) race. Each batch contains 4 external pooled QC samples (shown in black/dark colors) plus study samples distributed across demographic and clinical categories. The relatively uniform distribution of characteristics across batches confirms effective randomization of case-control pairs to batches.

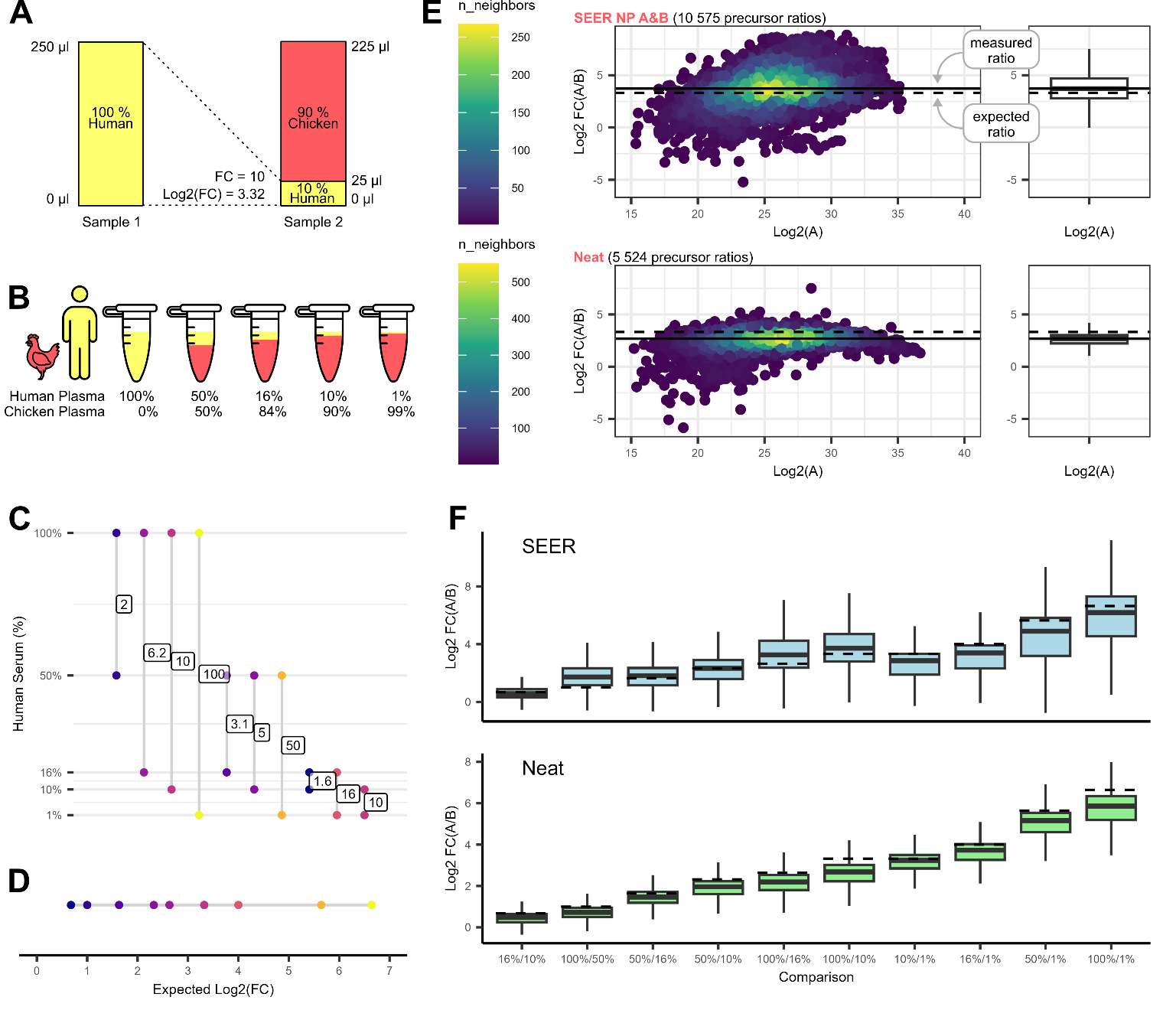

Figure S4: Mixed species verification of quantitative accuracy of proteomics pipeline. (A) Conceptual framework of the mixed-species assay. By comparing ratios of human proteins between mixtures, measured fold changes can be compared to expected fold changes across the measurable proteome. (B) Five different ratio mixtures, performed in experimental duplicate, were used to validate the workflow. (C) The ratio measurements enable quantification of human proteins across 10 different fold change comparisons ranging from 2-fold to 100-fold (color-coded by expected fold change). (D) Expected measurable fold changes plotted on a log₂ scale, with colors corresponding to the mixture compositions in panel C. (E) Representative MA-style plots comparing measured log₂ fold changes (FC(A/B)) versus mean precursor abundance (Log₂(A)) for Seer NP A&B (10,575 precursor ratios, top) and neat plasma (5,524 precursor ratios, bottom). Points are colored by local density (n_neighbors). Dashed horizontal lines indicate expected ratios; boxplots (right) show the distribution of measured ratios across all precursors. (F) Boxplots comparing median log₂ fold changes across different mixture comparisons for Seer (blue) and Neat (green) workflows. Both methods show measured fold changes that track closely with expected ratios across the dynamic range tested. Seer samples were processed with the Seer Proteograph XT assay, with equal masses of NP A and NP B peptides combined prior to analysis. Neat samples were processed using traditional urea denaturation and tryptic digestion. All samples were analyzed with 500 ng on-column loading using Evosep One (30 SPD) on an Exploris 480 operating in DIA mode.

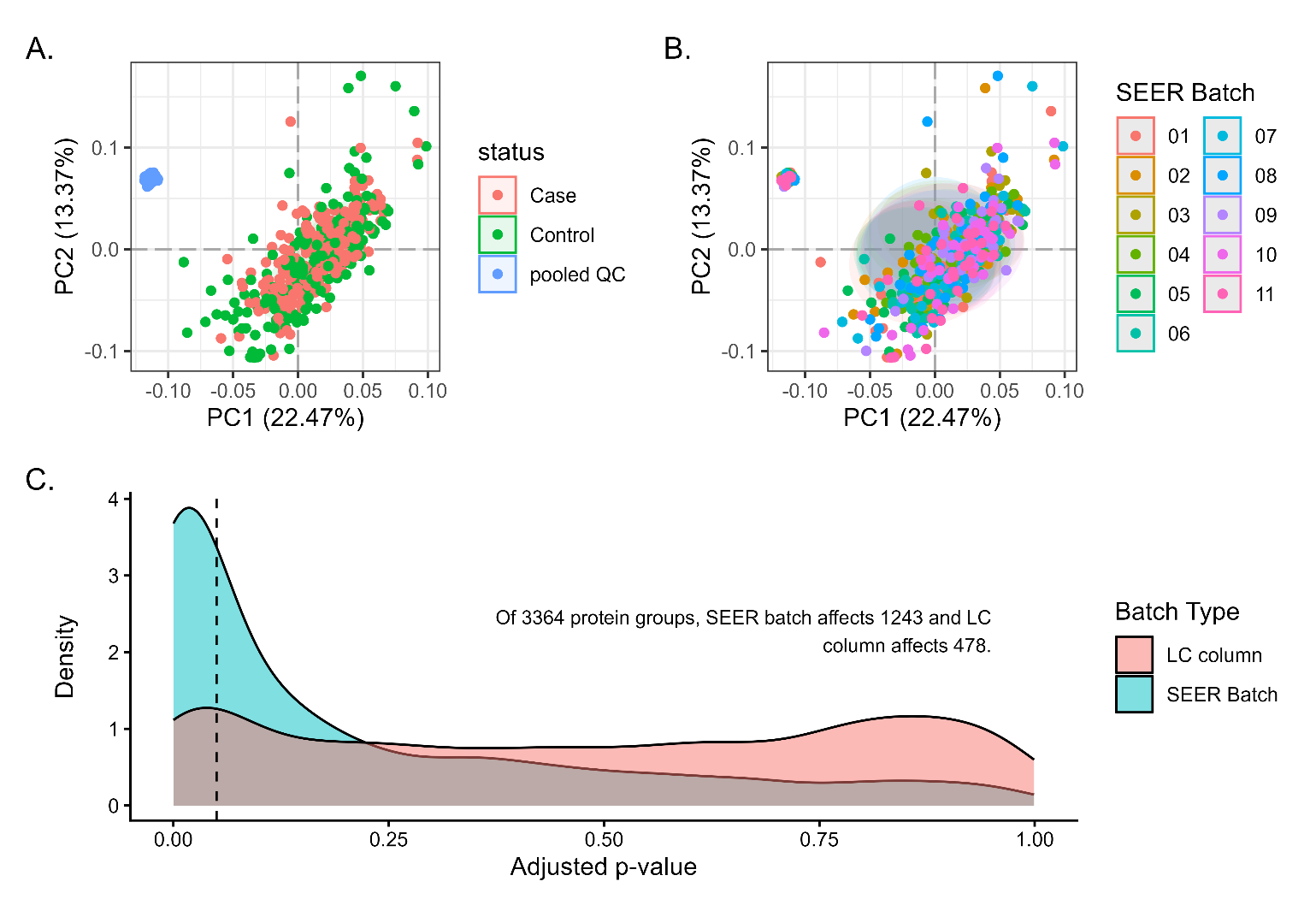

Figure S5: Seer batch is the primary driver of technical variation**.** PCA plots of uncorrected log₂ protein group intensities colored by **(A)** case-control status or **(B)** Seer batch. The tight clustering visible in panel A represents external pooled QC samples, which are distinct from study samples due to their commercial source. **(C)** Distribution of adjusted p-values from limma testing for batch effects shows that Seer batch (pink) significantly affects the majority of protein groups, while LC-MS column effects (teal) are minimal. Dashed line indicates adjusted p-value threshold of 0.05.
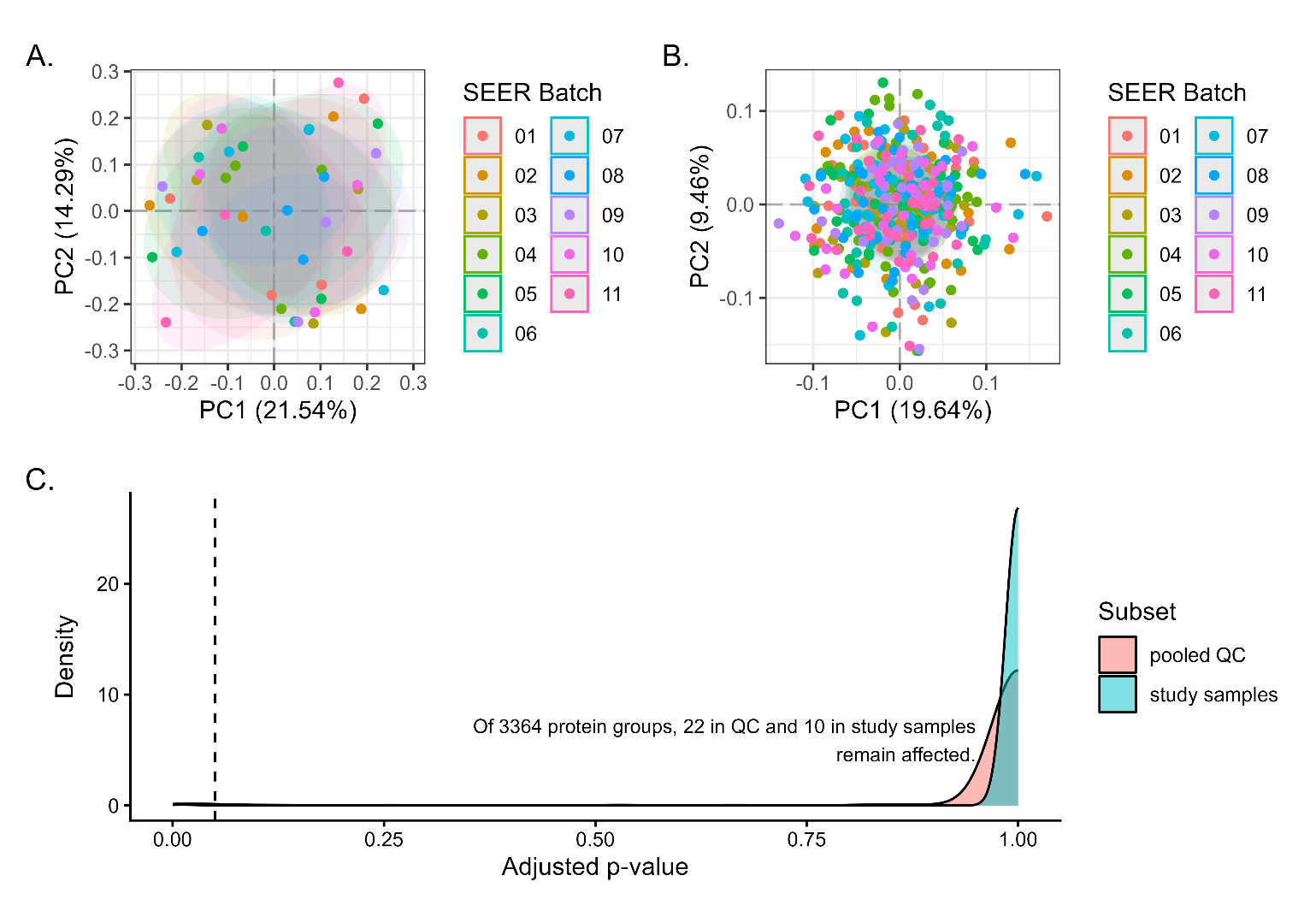

Figure S6: Separate batch correction strategy for pooled QC and study samples. Due to distinct variance patterns between external pooled QC serum and DODSR study specimens, datasets were separated and batch corrected independently using ComBat. (A) PCA plot of batch-corrected pooled QC samples colored by Seer batch, showing effective removal of batch effects. (B) PCA plot of batch-corrected study samples colored by Seer batch, demonstrating successful batch correction without compromising biological variation. (C**)** Distribution of adjusted p-values from limma testing for testing between batches for pooled QC samples (pink) and study samples (teal). After correction, few protein groups remained significantly affected by batch (22 in pooled QC, 10 in study samples out of 3,364 total protein groups), confirming effective batch correction in both subsets.

Table S1: Protein groups quantified in at least the given fraction of samples, for the whole study and for the samples entering each modeling comparison.

|  |  | Count of protein groups in fraction of samples | | | | |
| --- | --- | --- | --- | --- | --- | --- |
| Sample set | Samples (n) | 100% | ≥ 75% | ≥ 50% | ≥ 25% | ≥ 1 sample |
| All samples | 434 | 455 | 1424 | 1914 | 2474 | 3364 |
| All study samples | 390 | 463 | 1424 | 1914 | 2477 | 3364 |
| All pooled QC samples | 44 | 1059 | 1598 | 1946 | 2360 | 3111 |
| Cross-sectional: 2 yr prior | 132 | 568 | 1417 | 1887 | 2463 | 3325 |
| Cross-sectional: 4 yr prior | 106 | 526 | 1408 | 1901 | 2476 | 3351 |
| Cross-sectional: 2 yr post | 82 | 637 | 1445 | 1920 | 2430 | 3286 |
| Cross-sectional: DX window | 70 | 719 | 1512 | 1991 | 2533 | 3348 |
| Longitudinal: 4 yr prior → 2 yr prior | 200 | 494 | 1400 | 1882 | 2476 | 3359 |
| Longitudinal: 2 yr prior → DX window | 128 | 612 | 1467 | 1949 | 2532 | 3352 |
| Longitudinal: 4 yr prior → DX window | 100 | 584 | 1451 | 1929 | 2510 | 3353 |

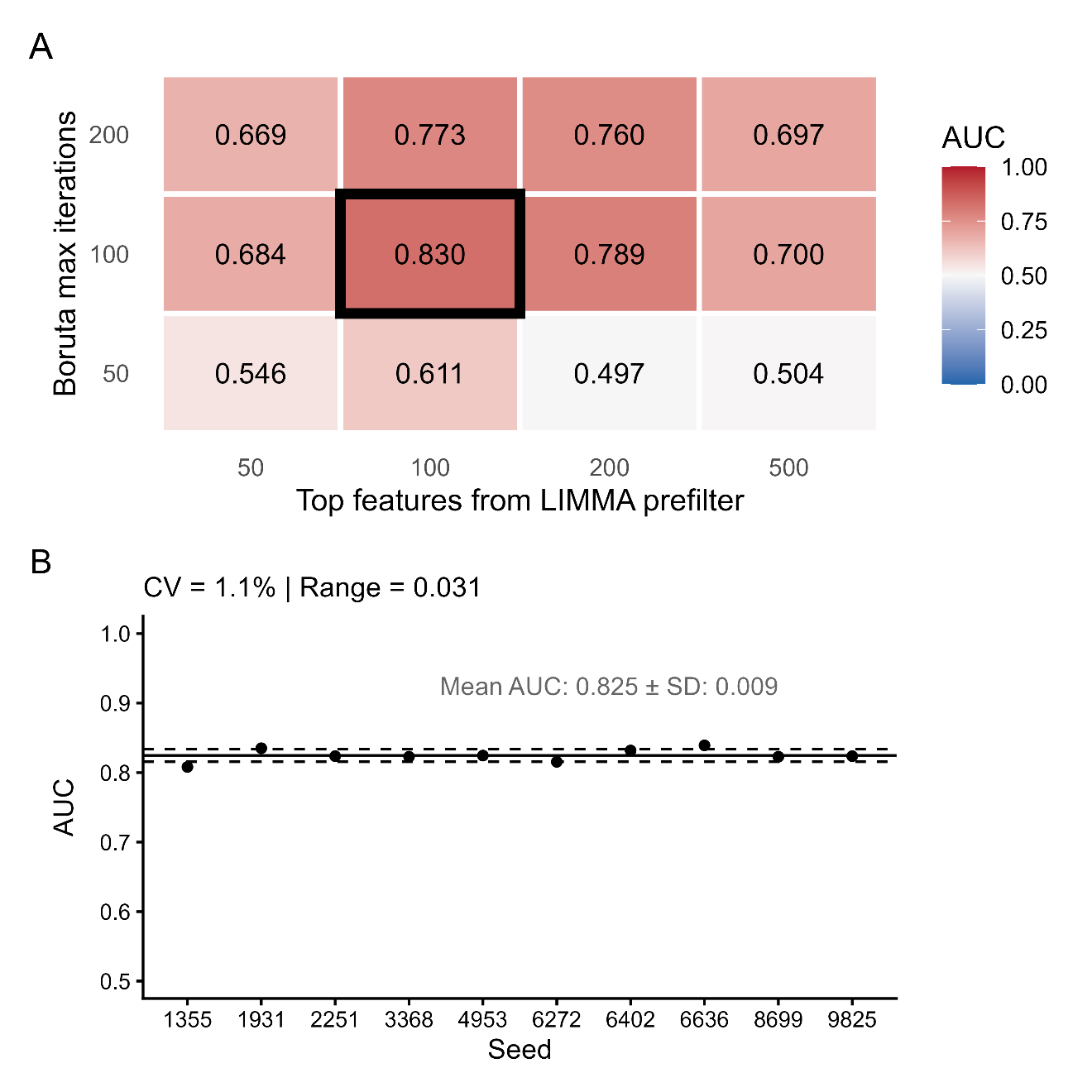

Figure S7: **Random forest model hyperparameter optimization and stability validation.** (A) Grid search AUC performance across limma prefilter sizes and Boruta iteration limits. Leave-one-out cross-validation results for DX window cross-sectional comparison (n=70; 35 cases, 35 controls) using 1,581 features (S/N ≥ 2, presence ≥ 50%). Values show mean AUC; optimal parameters indicated by box. (B) Model stability across random seeds at optimal hyperparameters (prefilter_n=100, boruta_maxRuns=100). Each point represents one LOOCV run, solid and dashed lines indicate mean and standard deviation, respectively.

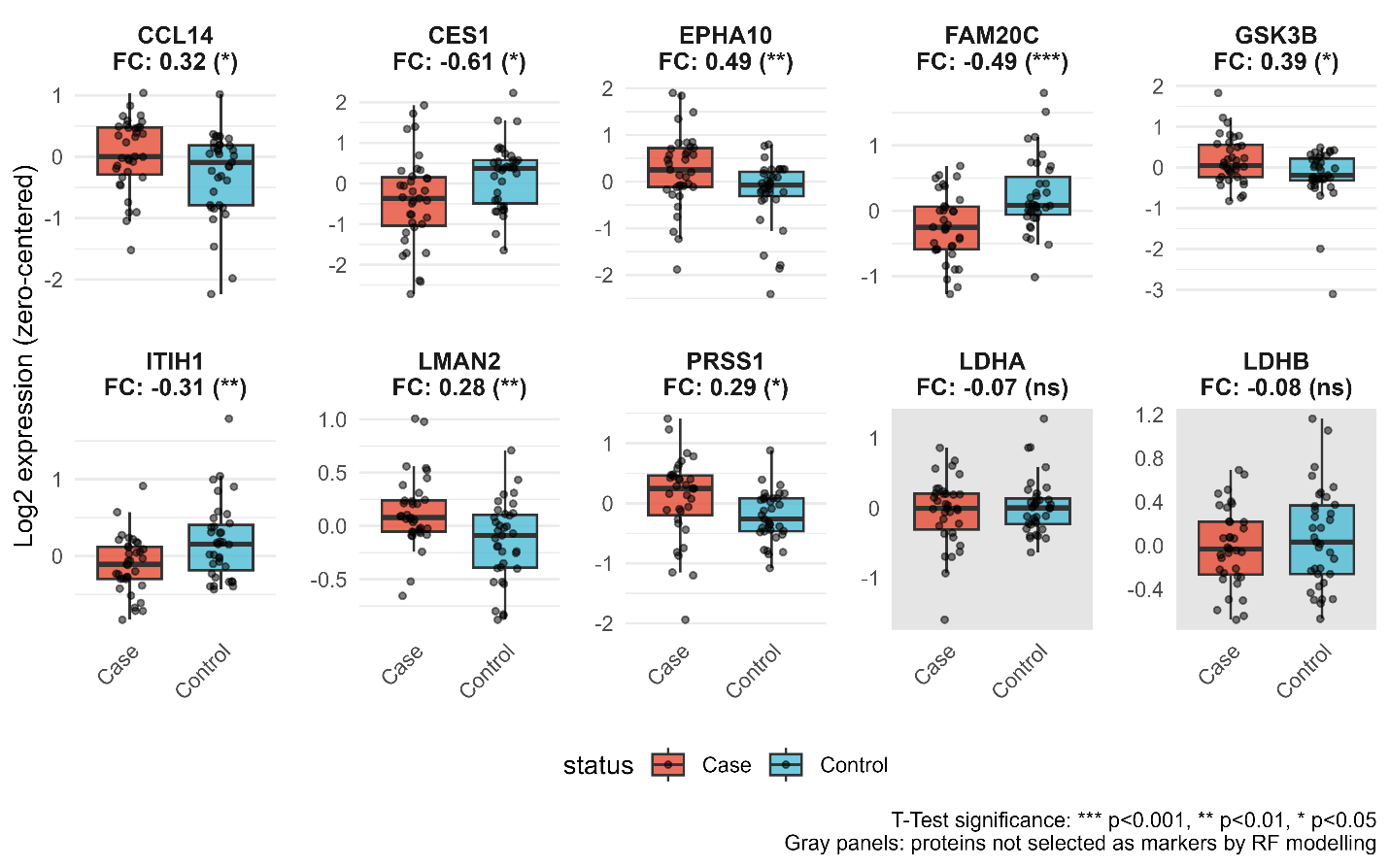

Figure S8: **Differential expression of consensus signature proteins and established melanoma markers in the diagnosis window (DX window).** Box plots show log2 expression (zero-centered) for the 8 consensus proteins selected in ≥50% of cross-validation folds (CCL14, CES1, EPHA10, FAM20C, GSK3B, ITIH1, LMAN2, PRSS1), plus lactate dehydrogenase subunits LDHA and LDHB for comparison. Fold change (FC) indicates case relative to control; positive values indicate higher expression in cases. LDHA and LDHB, components of the established advanced-stage melanoma marker LDH, were detected but did not significantly differentiate cases from controls, consistent with the early-stage composition of this cohort. T-test significance: *** p<0.001, ** p<0.01, * p<0.05, ns = not significant.

Table S2: Identification evidence per marker. Identification and quantification used the search database described in the Methods (GENCODE release 42 and Ensembl release 108); gene names and UniProt IDs are mapped from the Ensembl accessions for readability.

| Gene | UniProt  (mapped) | Ensembl protein group (searched) | Model | Peptides  identified | Study S/N^†^ | Evidence |
| --- | --- | --- | --- | --- | --- | --- |
| CCL14 | A0A0B4J2G5 | ENSP00000484818 | DX window | 1 | 4.9 | Single peptide |
| CES1 | P23141 | ENSP00000353720; ENSP00000355193; ENSP00000390492 | DX window | 17 | 68.8 | Multi-peptide |
| EPHA10 | Q5JZY3; J3KQG3 | ENSP00000316395; ENSP00000362139; ENSP00000397746 | DX window | 1 | 17.9 | Single peptide |
| FAM20C | Q8IXL6 | ENSP00000322323 | DX window | 6 | 16.8 | Multi-peptide |
| GSK3B | A0A3B3ITW1 | ENSP00000497956 | DX window | 5 | 5.6 | Multi-peptide |
| ITIH1 | P19827 | ENSP00000273283 | DX window | 47 | 35.1 | Multi-peptide |
| LMAN2 | Q12907 | ENSP00000303366 | DX window | 6 | 3.2 | Multi-peptide |
| PRSS1 | P07477;  E7EQ64 | ENSP00000308720; ENSP00000417854 | DX window + Longitudinal | 3 | 13.0 | Multi-peptide |
| PRSS2 | P07478; A6XMV9; A0A0J9YYC8 | ENSP00000488338; ENSP00000488437; ENSP00000488789 | Longitudinal | 4 | 45.5 | Multi-peptide |
| ST6GAL1 | P15907 | ENSP00000169298 | Longitudinal | 10 | 81.9 | Multi-peptide |
| VCAN | P13611 | ENSP00000265077 | Longitudinal | 33 | 31.3 | Multi-peptide |
| ^†^ See Eq. 1 in main text for definition of Study S/N. | | | | | | |

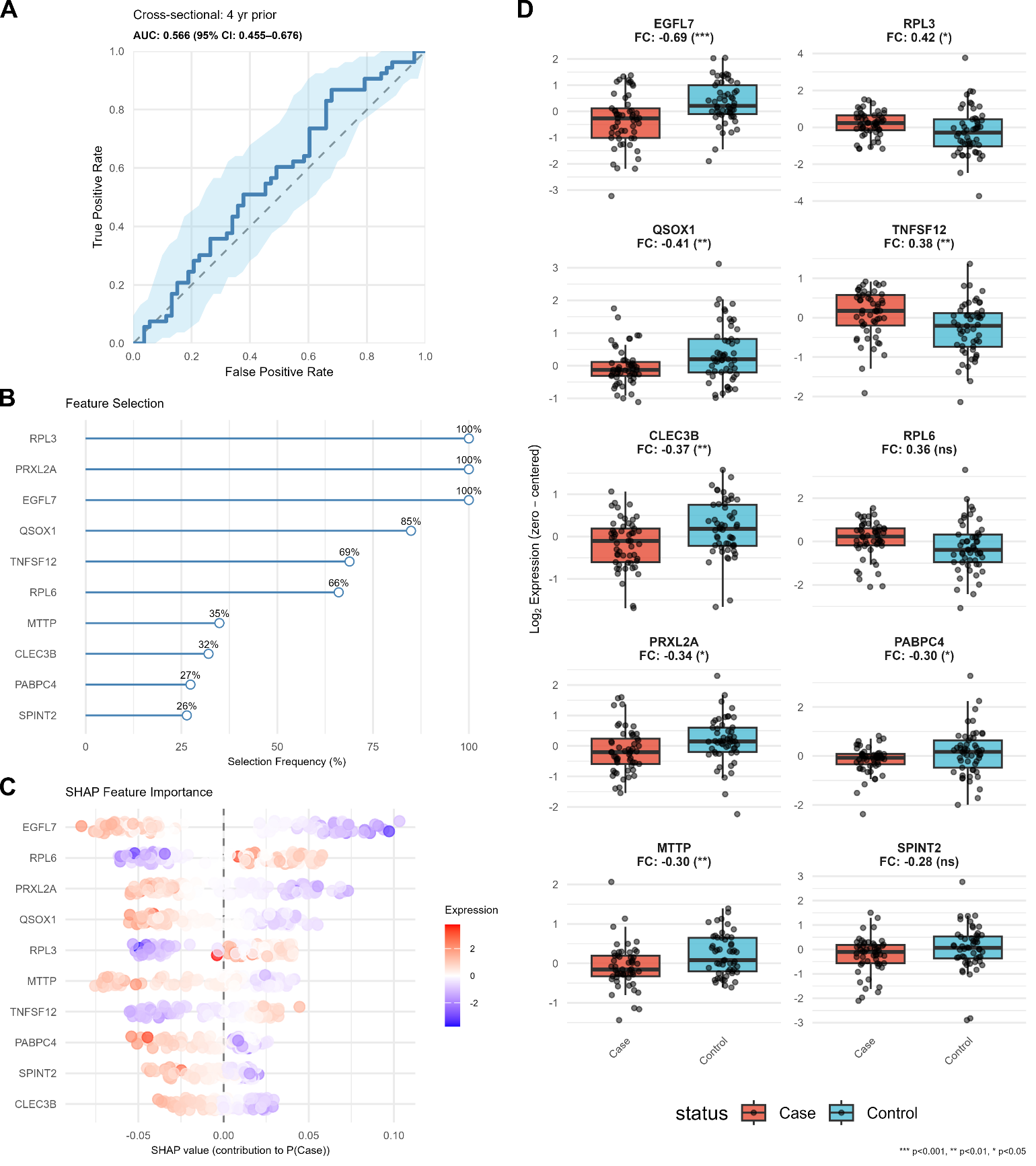

Figure S9**: LOOCV random forest classification for case vs. control at 4 years prior to diagnosis**. (A) ROC curve with 95% CI. (B) Boruta feature selection frequency (≥25% of folds shown). (C) SHAP beeswarm plot from consensus model (blue = low expression, red = high). (D) Case–control boxplots with log₂ fold change and t-test significance.

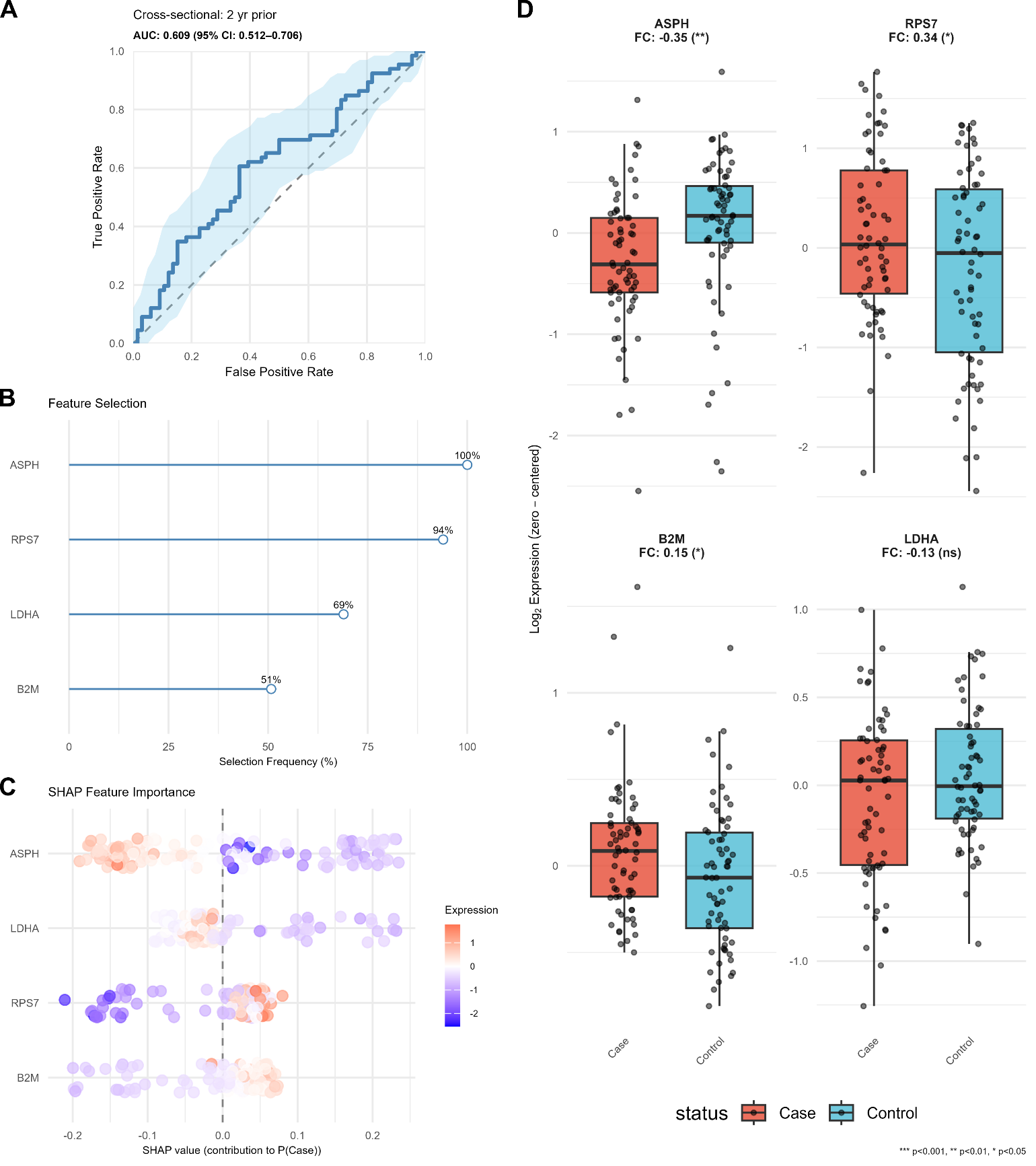

Figure S10: **LOOCV random forest classification for case vs. control at 2 years prior to diagnosis.** (A) ROC curve with 95% CI. (B) Boruta feature selection frequency (≥25% of folds shown). (C) SHAP beeswarm plot from consensus model (blue = low expression, red = high). (D) Case–control boxplots with log₂ fold change and t-test significance.

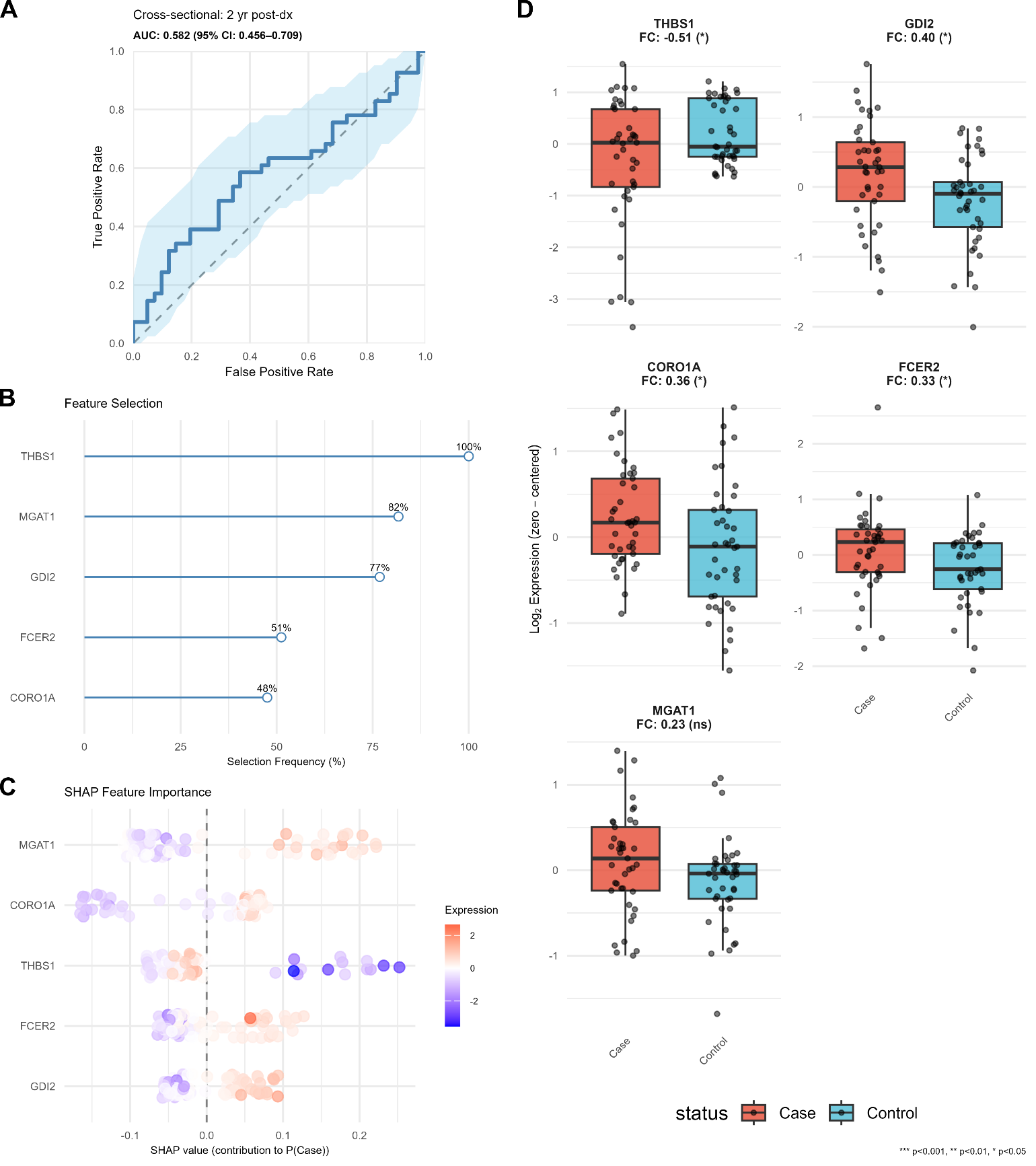

Figure S11**: LOOCV random forest classification for case vs. control at 2 years post-diagnosis.** (A) ROC curve with 95% CI. (B) Boruta feature selection frequency (≥25% of folds shown). (C) SHAP beeswarm plot from consensus model (blue = low expression, red = high). (D) Case–control boxplots with log₂ fold change and t-test significance.

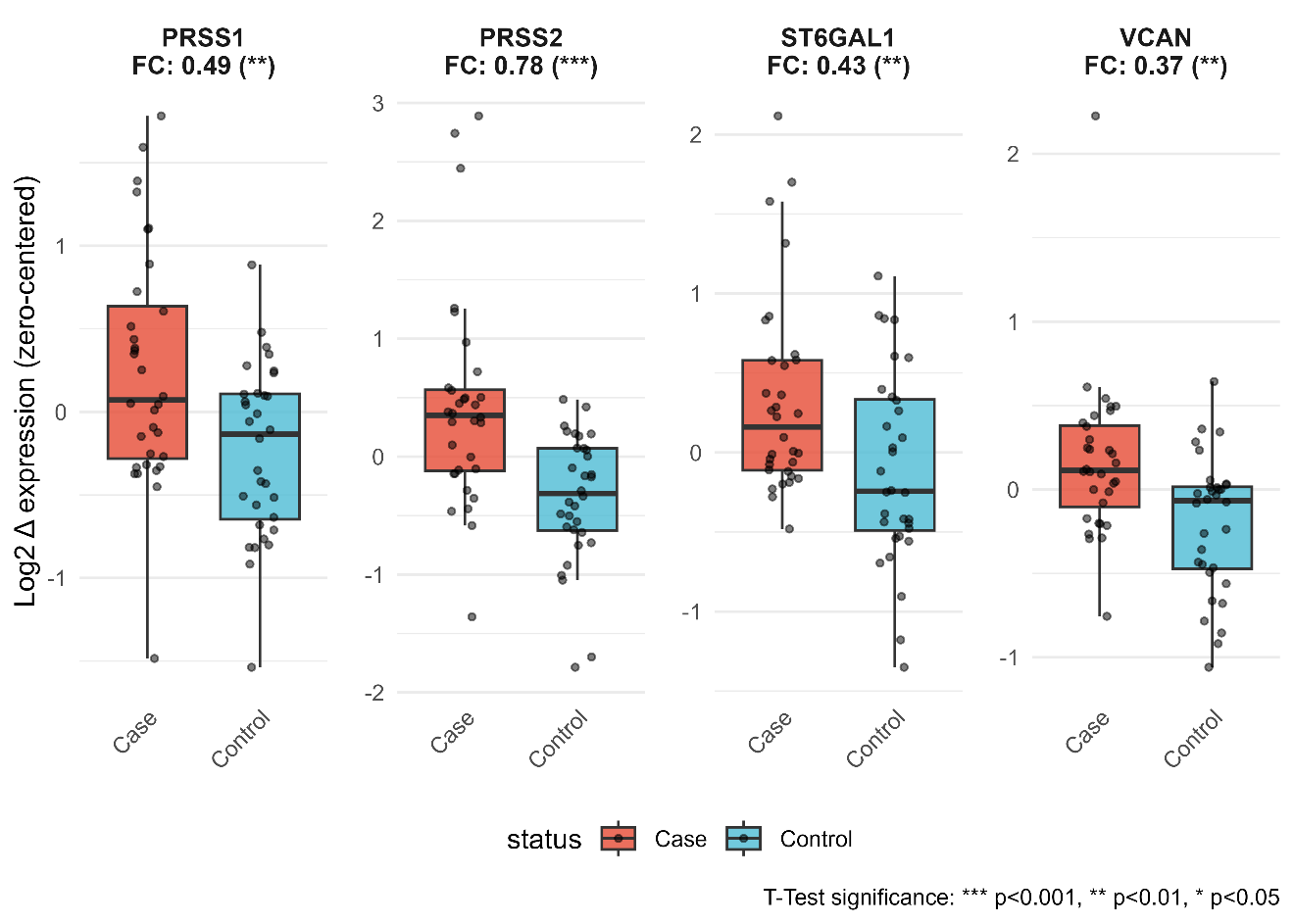

Figure S12: **Univariate comparison of top RF-selected candidate markers for longitudinal protein expression changes between the diagnosis window and 2 years prior.** Log₂ protein expression changes (batch-corrected, zero-centered) for the 4 features selected in ≥ 50% of LOOCV folds for DX window – 2 yr prior. Each panel shows case-control comparisons (n = 32 per group) with fold change (FC) values and t-test statistical significance (unadjusted p-values: *** p<0.001, ** p<0.01, * p<0.05). Raw p-values are reported as these represent univariate comparisons of RF-selected features on the same samples used for feature selection, not proteome-wide screening or independent validation.

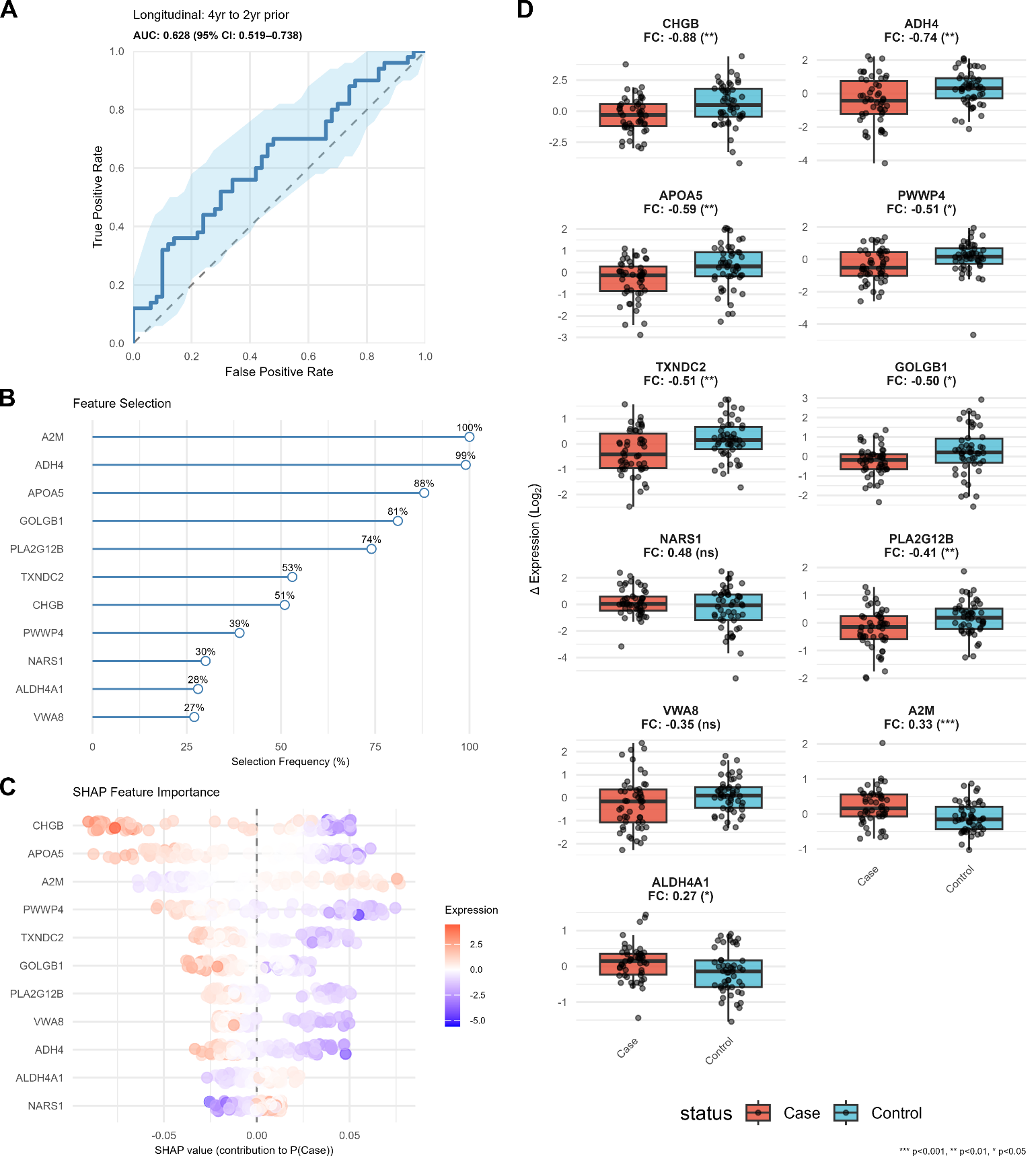

Figure S13**: LOOCV random forest classification of Δ expression (2 yr prior − 4 yr prior) between cases and controls.** (A) ROC curve with 95% CI. (B) Boruta feature selection frequency (≥25% of folds shown). (C) SHAP beeswarm plot from consensus model (blue = low Δ expression, red = high). (D) Case–control boxplots of Δ expression with fold change and t-test significance.

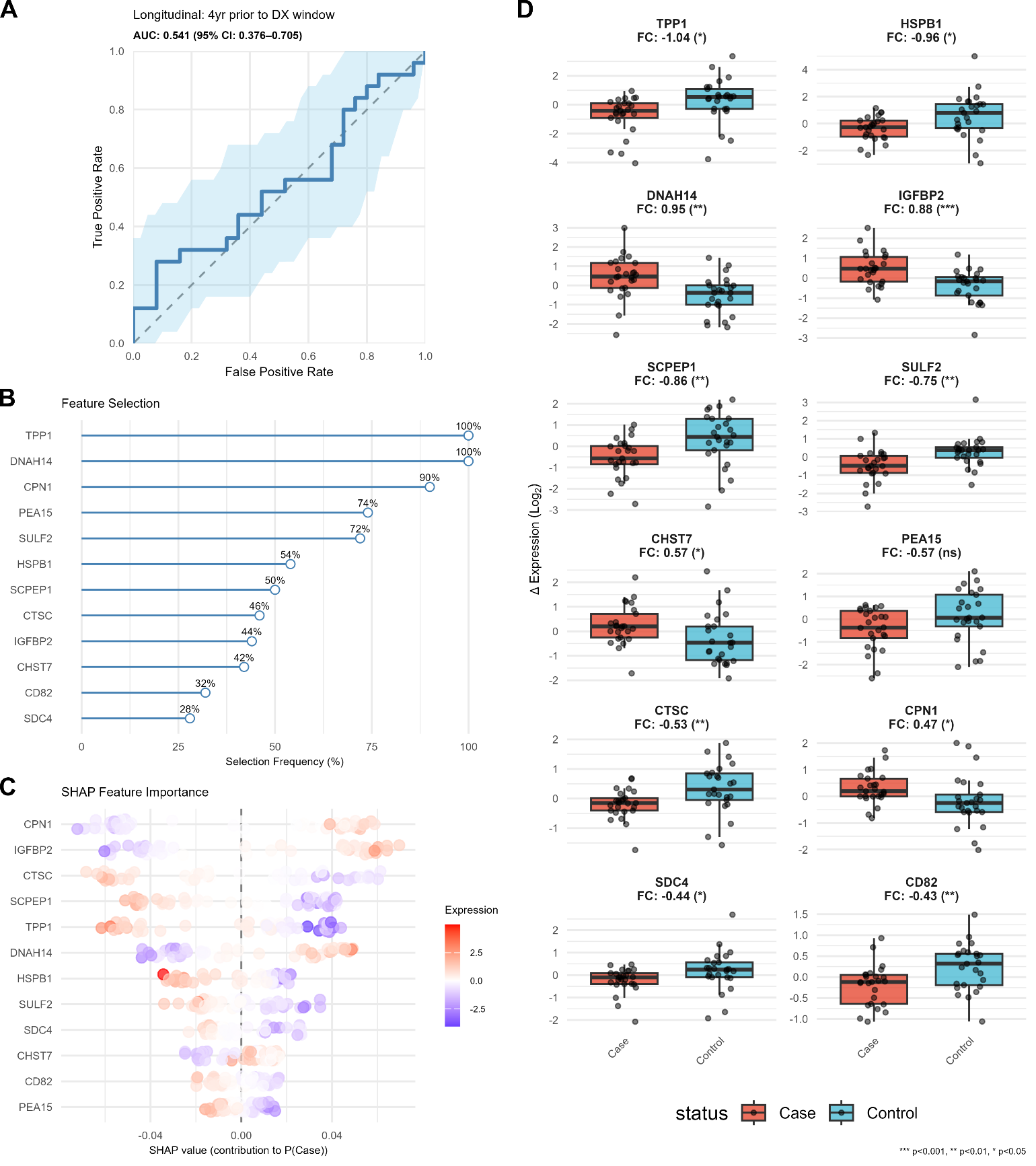

Figure S14: **LOOCV random forest classification of Δ expression (DX window − 4 yr prior) between cases and controls.** (A) ROC curve with 95% CI. (B) Boruta feature selection frequency (≥25% of folds shown). (C) SHAP beeswarm plot from consensus model (blue = low Δ expression, red = high). (D) Case–control boxplots of Δ expression with fold change and t-test significance.

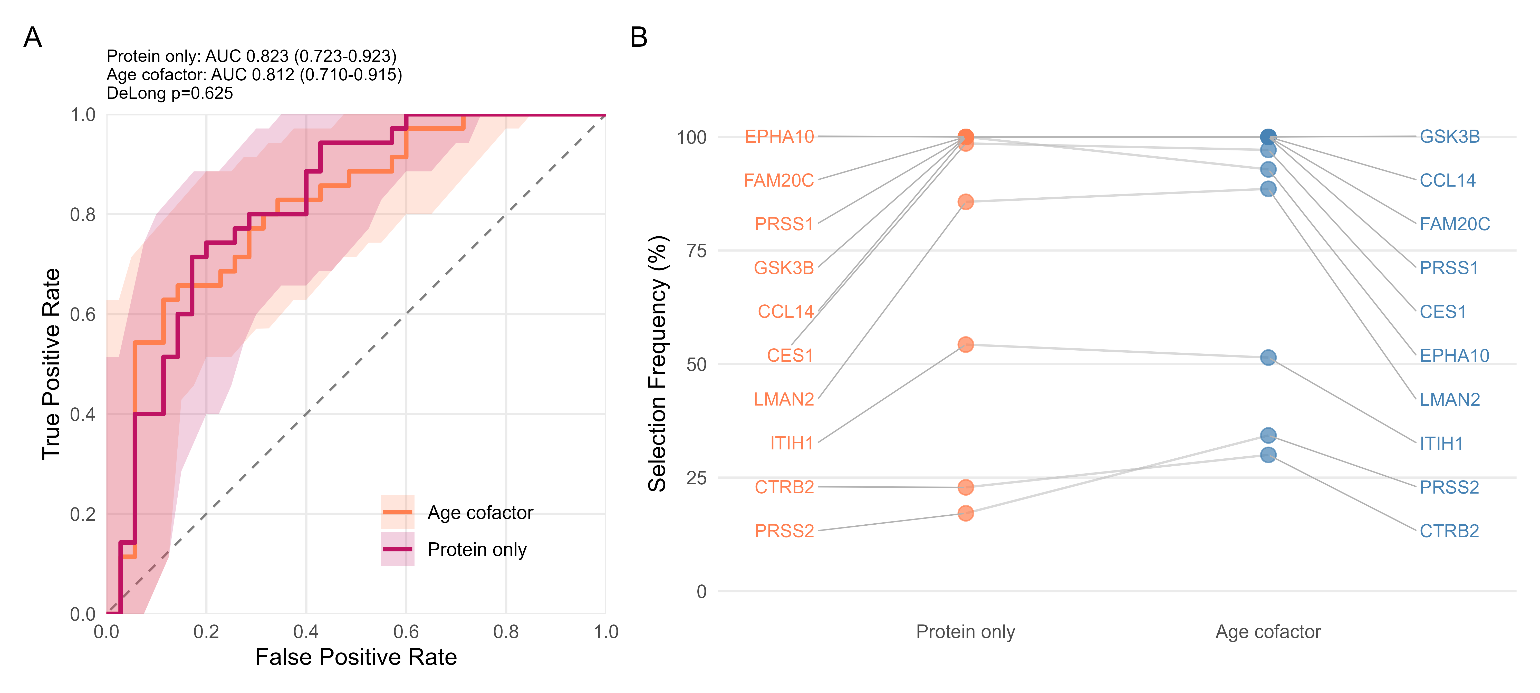

Figure S15: **Addition of age as a cofactor does not improve model performance in the diagnosis window.** **(A)** ROC curves comparing random forest models trained with protein features only (pink, AUC=0.823, n=70) versus protein features plus subject age as a cofactor (orange, AUC=0.812, n=70) for the DX window cross-sectional comparison. Shaded regions represent 95% confidence intervals. The two models' AUCs were not significantly different (DeLong test(1), p = 0.625). **(B)** Comparison of feature selection frequency across leave-one-out cross-validation folds between the two modeling approaches. Lines connect the same protein across models, showing how selection frequency changed when age was included.

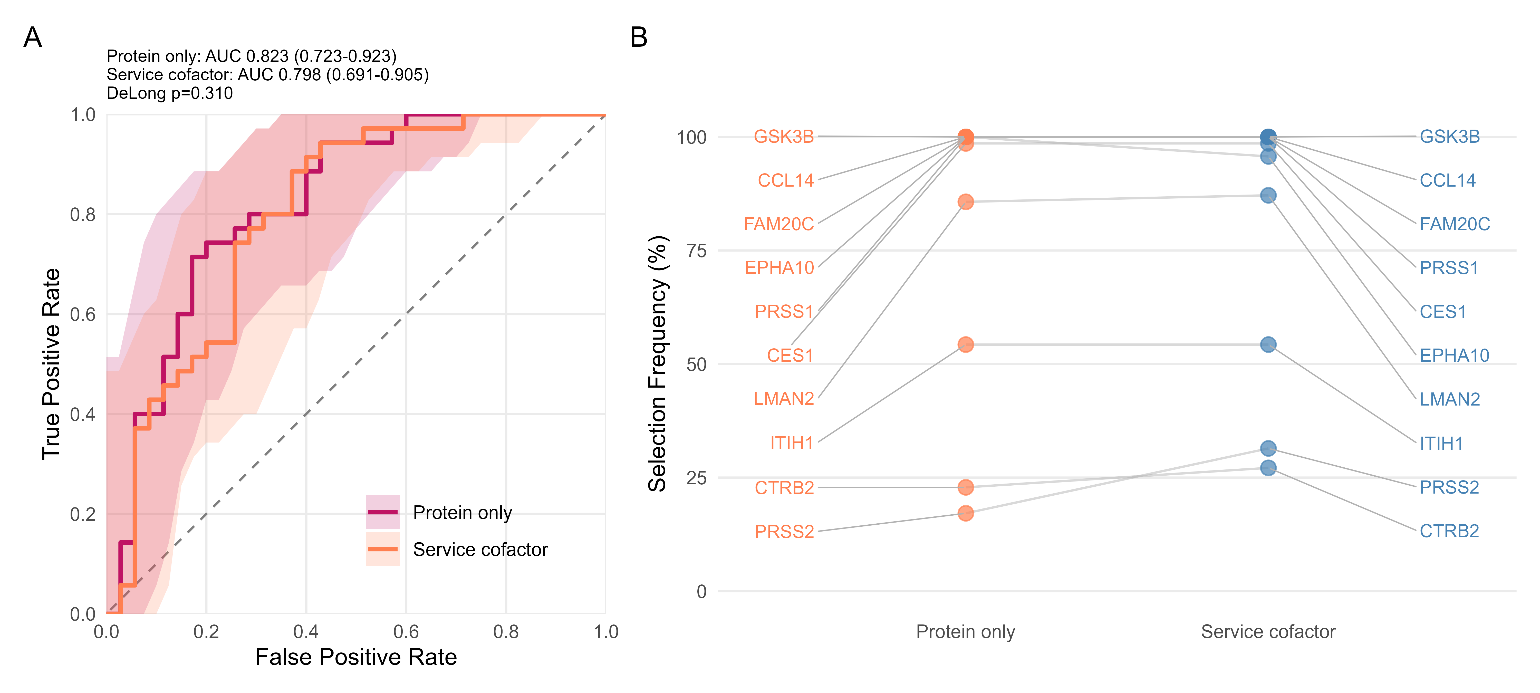

Figure S16**: Addition of service branch as a cofactor does not improve model performance in the diagnosis window.** **(A)** ROC curves comparing random forest models trained with protein features only (pink, AUC=0.823, n=70) versus protein features plus subject service branch at time of diagnosis as a cofactor (orange, AUC=0.798, n=70) for the DX window cross-sectional comparison. Shaded regions represent 95% confidence intervals. The two models' AUCs were not significantly different (DeLong test(1), p = 0.310). **(B)** Comparison of feature selection frequency across leave-one-out cross-validation folds between the two modeling approaches. Lines connect the same protein across models, showing how selection frequency changed when service branch was included.

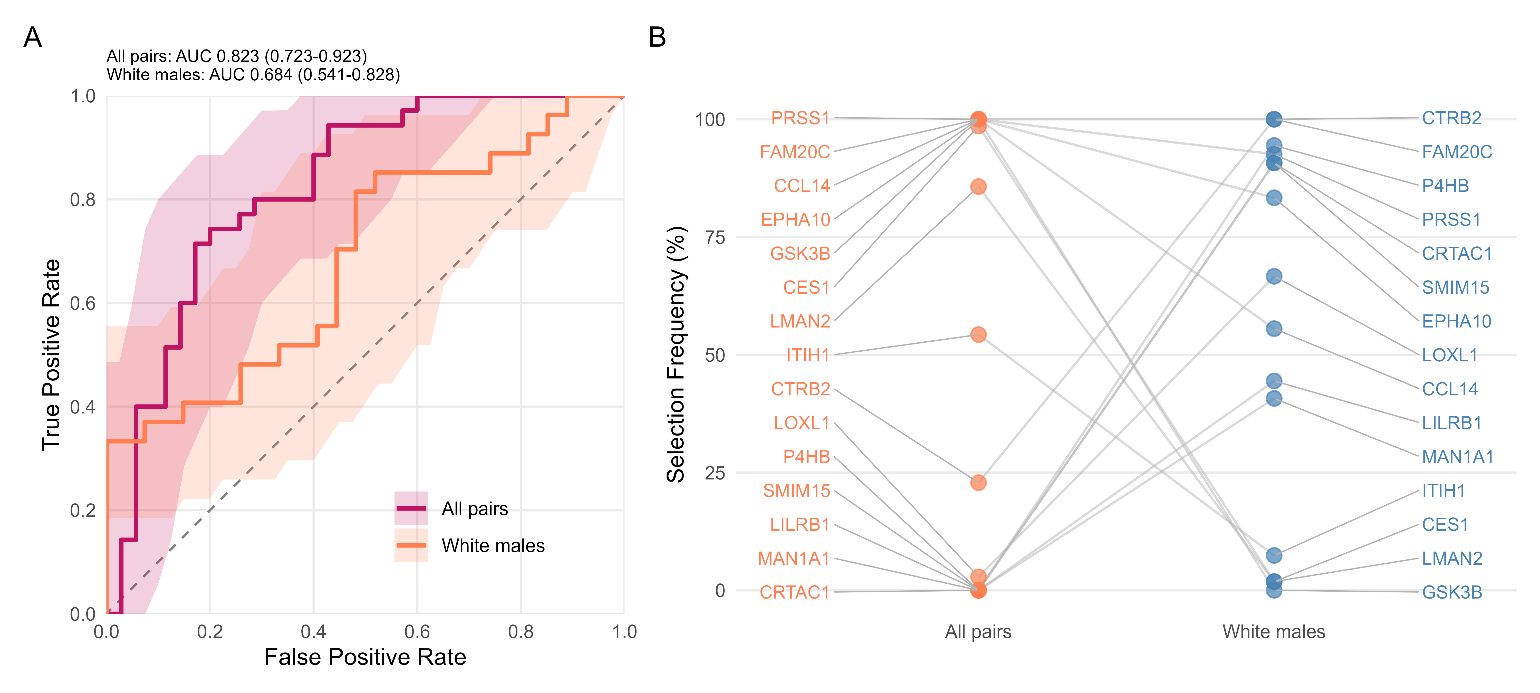

Figure S17: **Demographic subsetting to white males reduces model performance and cohort size.** **(A)** ROC curves comparing random forest models for all case-control pairs (pink, AUC=0.823, n=70) versus white males only (orange, AUC=0.684, n=54) at the DX window timepoint. Shaded regions represent 95% confidence intervals. **(B)** Comparison of feature selection frequency across leave-one-out cross-validation folds. Lines connect the same protein across the two cohorts. Note DeLong test not applicable as models were evaluated on different sample sizes.

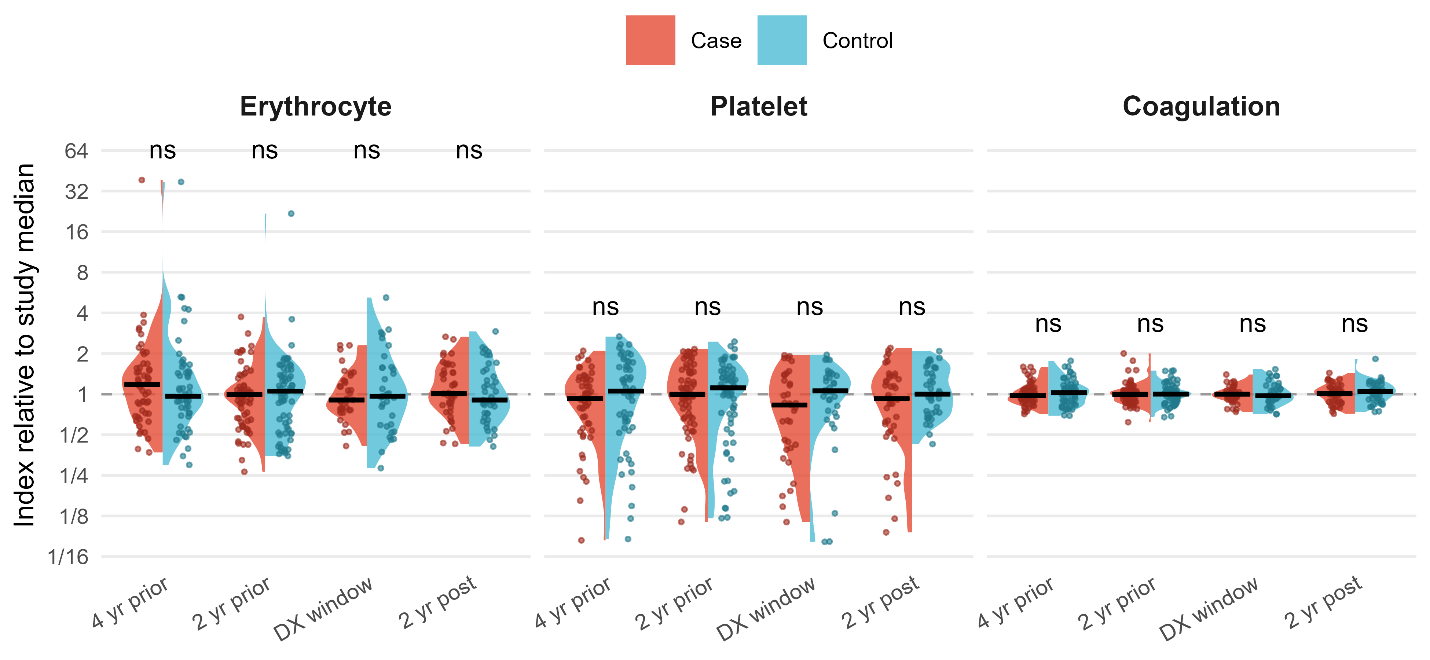

Figure S18**:** **Sample-quality marker panels in cases and matched controls**. Erythrocyte and platelet indices: summed intensity of the erythrocyte or platelet marker panel of Geyer et al. (2) as a fraction of total protein intensity. Coagulation index: total protein intensity divided by the summed intensity of the coagulation proteins depleted by clotting (2). Indices are shown relative to their median across all study samples (dashed line); black bars indicate group medians. Cases and matched controls were compared at each timepoint by paired t-test on log2 indices; ns = not significant (p > 0.05). Geyer et al. defined the platelet panel in plasma, where it measures residual platelets left after centrifugation. In serum, platelets are activated and largely retained in the clot, so the index reflects platelet-derived protein carried into the serum; poor clot formation would therefore appear as values above the median, not as the low tail. Panel proteins quantified (detected/defined by Geyer et al.): erythrocyte 27/29, platelet 28/29, coagulation 12/12, each detected in a median of 100% of study samples.

### References

1.	DeLong ER, DeLong DM, Clarke-Pearson DL. Comparing the areas under two or more correlated receiver operating characteristic curves: a nonparametric approach. Biometrics. 1988;44:837–45.

2.	Geyer PE, Voytik E, Treit PV, Doll S, Kleinhempel A, Niu L, et al. Plasma Proteome Profiling to detect and avoid sample-related biases in biomarker studies. EMBO Mol Med. 2019;11:e10427. https://doi.org/10.15252/emmm.201910427
